# Mapping long-term memory modulation with transcranial alternating current stimulation

**DOI:** 10.1101/2025.10.27.684796

**Authors:** Dima Chitic, Miles Wischnewski

## Abstract

**Background:** Long-term memory (LTM), which is the ability to retain information over a long period of time, is governed by neural oscillations in various parts of the cerebral cortex. To investigate the relationship between oscillations and memory performance, transcranial alternating current stimulation (tACS) has been used to modulate LTM. However, due to the variability in stimulation parameters and locations, no clear picture has arisen on where tACS needs to be applied to improve LTM.

**Methods:** We collected behavioral data from 20 studies (58 effect sizes) and computationally modeled the corresponding brain electric fields based on each study’s unique tACS parameters. As a result, we generated a map of performance–electric field index (PEI) values using linear and quadratic fits.

**Results:** Theta-tACS showed no regions with a positive linear association with LTM. However, left lateral prefrontal and temporal areas exhibited negative associations, meaning that theta stimulation of that region led to decreased LTM performance. A quadratic analysis revealed a U-shaped relation in the medial temporal and ventromedial prefrontal cortex, suggesting improved memory at low and high, but not moderate, electric field strength values. Gamma-tACS showed positive linear associations in parieto-occipital cortex and negative ones in frontal regions. For gamma, we found no regions that expressed a quadratic relationship.

**Conclusion:** These findings indicate that LTM-relevant tACS effects are frequency- and site-specific and potentially nonlinear with respect to intensity. Integrating meta-analytic effect sizes with individualized field estimates provides a brain-wide framework for identifying targets of memory modulation and for guiding future fundamental and translational research.

## Introduction

The ability to retain information from hours, days, weeks, or even months ago is one of the hallmarks of the brain^1^. Besides the importance of gaining a fundamental understanding of the underlying brain mechanisms, long-term memory (LTM) studies are critical for further expanding the treatment options for memory-related deficits, such as Alzheimer’s disease or traumatic brain injury^2,3^. In rodent models, memory is most commonly studied in subcortical areas, including the hippocampus and the amygdala^4–6^. However, human studies tend to focus on various cortical regions involved in processing and retaining information, including prefrontal, parietal, and temporal structures ^7–12^. Furthermore, within these regions, particular neural oscillations, such as theta (3-8 Hz) and gamma (30-200 Hz), have been highlighted through their association with encoding and retrieval of memory^13–15^.

Previous studies have investigated the role of oscillations using neurophysiological recordings. Osipova et al^16^ engaged their participants in a picture recognition task while magnetoencephalographic (MEG) data were recorded. They found higher theta and gamma power both during encoding when the picture was subsequently remembered (subsequent memory effect) and during retrieval, when distinguishing old pictures from new. Furthermore, intracranial recordings from epilepsy patients showed a similar pattern, with increases in theta and gamma power over several cortical regions for subsequent memory effects^17–19^. In contrast, Hanouneh et al^20^ showed that theta power in prefrontal regions was negatively correlated with LTM retrieval. However, Scholz et al^21^ showed that theta power is positively associated with memory encoding.

Given the correlational nature and variability in results of the studies using electrophysiological techniques, others have used neuromodulation methods to pinpoint regions that causally relate to memory performance^22–24^. One particular technique suited to study neural rhythms is transcranial alternating current stimulation (tACS) and various studies have used it to alter cortical oscillations during memory tasks^25–28^. By applying a sinusoidal low-intensity current to the scalp, tACS has been shown to entrain neural spiking, which can result in a modulation of endogenous oscillatory activity^29,30^. Besides these neurophysiological effects, various studies have shown that tACS is able to modulate human behavior and cognition^25,31,32^. The rationale is that the tACS-induced entrainment of neural oscillations is directly related to changes in behavioral performance. As an example, long-term face recognition was improved by applying theta tACS to the right posterior parietal region during encoding^33^. On the other hand, Manippa et al^26^ found reduced LTM performance following gamma tACS over bilateral temporal areas.

In accordance with the breadth of results in MRI and EEG studies, tACS has been applied in LTM experiments at various cortical locations and frequencies. Given the variety in parameters and protocols, a classic meta-analytic approach is suboptimal to summarize the efficacy of tACS, nor to identify core functional regions of LTM. By integrating meta-analytic results with electric field (E-field) modeling, this limitation can be overcome^34–37^. Using SimNIBS software, tACS-induced E-fields can be estimated based on a particular stimulation location and intensity^38^. By correlating the E-field distributions of different studies with their corresponding behavioral effect sizes (on LTM in this case), a meta-analytic brain map is generated that displays where tACS leads to increased, decreased, or no change in task performance^35^.

In the present study, we leverage the meta-modeling approach to summarize the effects of tACS on LTM. Thereby, we identify regions where neural oscillations in a given frequency are directly related to memory performance. Altogether, these results aim to complement previous research on cortical oscillations in LTM and point out cortical regions of interest for LTM research.

## Methods

### Sample

We collected effect sizes, expressed in Hedges’ g from available literature. The data was obtained from articles published in peer-reviewed journals, pre-prints, and published theses, by searching through PubMed, Web of Science, ProQuest (Dissertations and Theses), BioRxiv and PsycINFO databases. The search was conducted in general accordance with the PRISMA guidelines^39^. First, articles were identified using the following search terms: ("tACS" OR "transcranial alternating current stimulation") AND ("memory"). After removal of duplicates, the texts underwent abstract and full-text screening. Inclusion of studies in our modeling is based on the following criteria: I) Articles are written in English, II) Full-text is available to the researchers, III) manuscripts allowed for extraction of effect size (based on mean and standard deviation) from text, tables, or figures, IV) the study used a control condition (e.g., placebo) to which active stimulation can be compared, V) the study was conducted on a healthy adult population (>= 18 years old), VI) tACS applied online or offline, but not during sleep, VII) long-term memory assessed explicitly. For the criteria, we rely on a broad definition of long-term memory as information retrieved from the past that does not rely on active maintenance^2^. A preliminary search suggested that 15-25 articles could be eligible, suggesting that enough data can be collected to perform our meta-modeling analyses. In total, the final search (Figure 1) yielded 20 articles and 58 unique effect sizes (note that one study typically has multiple outcome measures). Across all studies n = 1210 healthy volunteers were included between the ages of 18-77. Since previously published data were used, no ethical approval was obtained.

**Figure 1.**
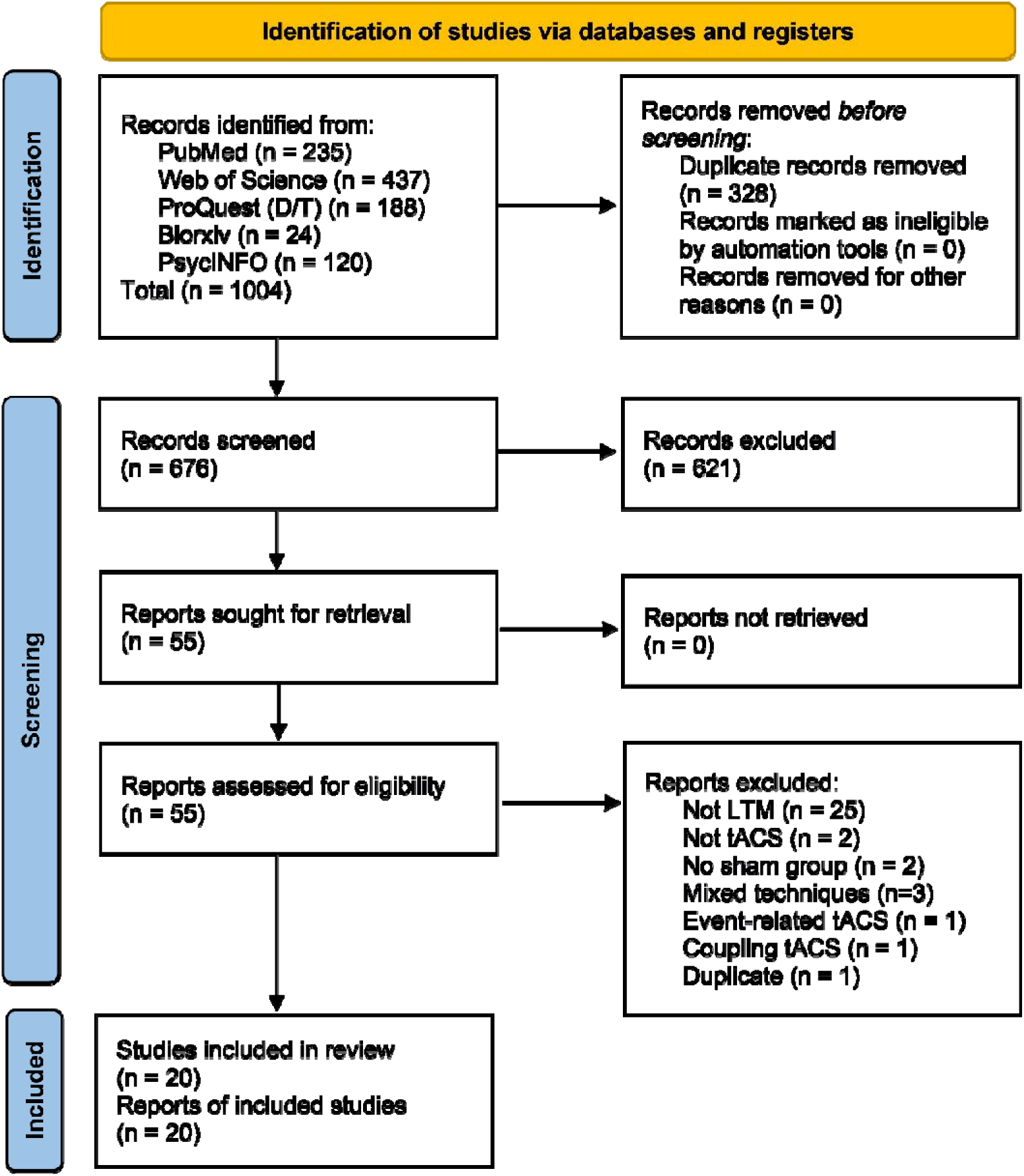
PRISMA overview of included studies.

### Effect size calculation

The standardized effect sizes were expressed as Hedges’ g values^40^. Hedges’ g values express the ratio between differences in effects and the variability, similar to Cohen’s d values^41^, with the addition of a correction factor for small sample sizes. If Hedges’ g values were reported, they were directly taken from the article. Alternatively, if the article provided the Cohen’s d value, the correction factor was applied to calculate Hedges’ g. If no effect size measures were directly reported, the Hedges’ g was calculated based on reported averages and standard deviations or those estimated from graphs. All effect sizes used in this analysis reflect the difference between an active stimulation condition and a sham control. As such, positive g values reflect improved memory performance in active tACS conditions compared to sham, while negative g values indicate worsened memory performance in active tACS conditions compared to sham.

### Electric field modeling

E-field simulations were performed on 50 realistic head models (age range 22-35, 22 females). The head models were generated based on data from the Human Connectome Project (HCP) S1200^42^. Quality and suitability for E-field modeling were assessed previously^43^. Each head model consists of 6 biological tissues, namely the skin, skull, cerebrospinal fluid, eyes, gray matter and white matter. For each tissue standard conductivity values were used (σskull□=□0.01□S/m, σskin□=□0.465 S/m, σCSF□=□1.654□S/m, σgraymatter□=□0.275□S/m, and σwhitematter□=□0.126□S/m)^44,45^. Electrode location, orientation, and material were gathered from the articles and used for E-field modeling in SimNIBS 4.1^38^. For the purpose of this study, we extracted the induced E-fields from the gray matter, as this reflects the tissue at which tACS most likely induces functional changes^46,47^. In order to compare and average between head models, all E-field simulations were transformed into the normalized FreeSurver average (fsavg) standard brain. Importantly, by first performing simulations in the individual head models and then transforming them into fsavg space (rather than directly performing electric-field simulations in fsavg models), individual variability is still accounted for.

### Calculation of the performance-electric field index

To relate different elements of fsavg model to functional brain regions, each element was categorized following the Human Connectome Project multimodal parcellation (HCP-MP) atlas^48,49^. This atlas parcellates the brain models into 360 function regions (180 per hemisphere). Given that we found 36 and 22 outcome measures for theta and gamma, respectively, and 360 functional regions, a 36 x 360 matrix and a 22 x 360 matrix of E-field values are obtained. These matrices are subsequently correlated with a vector of Hedges’ g values for each row of the matrix (i.e., for each functional region), resulting in two 1 x 360 vectors that represents the correlation between E-fields and tACS-induced performance changes per frequency^34–37^. We refer to these values as the performance-electric field index (PEI). Besides this, we report the uncorrected p-values. Note that due to the nature of meta-analytic datasets, no a priori power analysis was performed, and the number of data points depends on the available literature. As such, the exact p-value, and hence the optimal way to correct for multiple comparisons, is somewhat arbitrary. Therefore, we opted to report uncorrected p-values and visually represent them as p<0.05 (*), p<0.01 (**), and p<0.001 (***). For visualization, we categorized the 360 HCP-MP atlas region into 22 larger regions^50^.

In addition to the linear correlation analysis, we performed an exploratory analysis testing for a potential quadratic relationship between E-field strength and performance. This analysis was performed as several studies have shown that the effects of tACS follow a U-shape^51–53^ or an inverted U-shape ^54,55^. For each cortical region, we fit a linear (y = β_0_ + β_1_x) and a quadratic regression model (y = β₀ + β₁x + β₂x²), with the behavioral effect size (Hedges’ g) as the dependent variable and regional E-field magnitude as the predictor. For both linear and quadratic models, a correlation value was defined as the signed square root of R-squared, with the sign determined by the regression values being positive or negative. For each region, we determined the better-fitting model (linear or quadratic) based on adjusted R-squared. Across 50 head models, results were averaged per region. If more than 50% of the head models yielded a better fit for a quadratic regression in a particular region, the quadratic correlation values were averaged. We refer to this value as the PEI_QUAD,_ where positive and negative values correspond to a U-shaped and an inverted U-shaped relationship between E-field strength and performance, respectively.

**Table 1.**
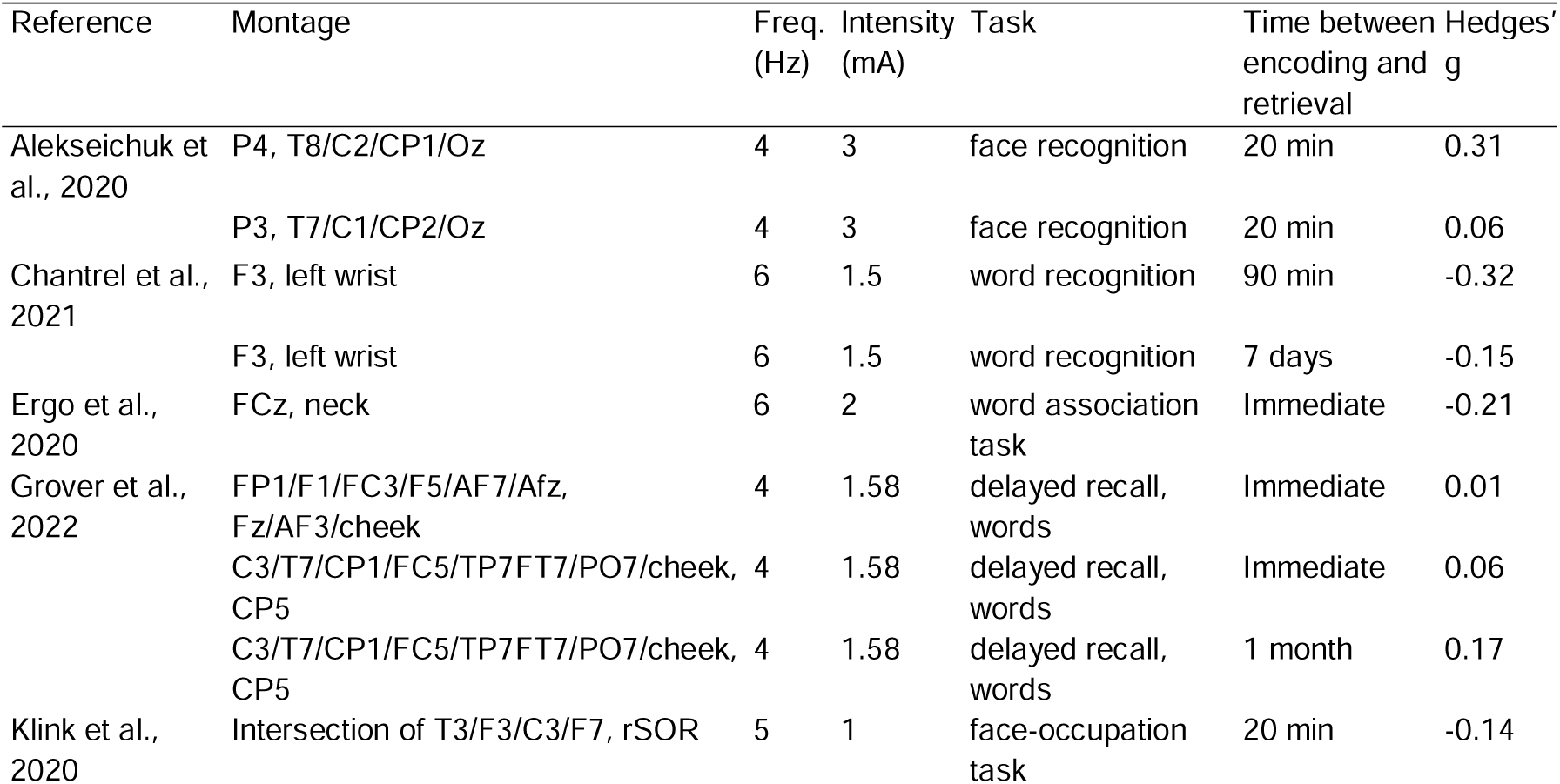
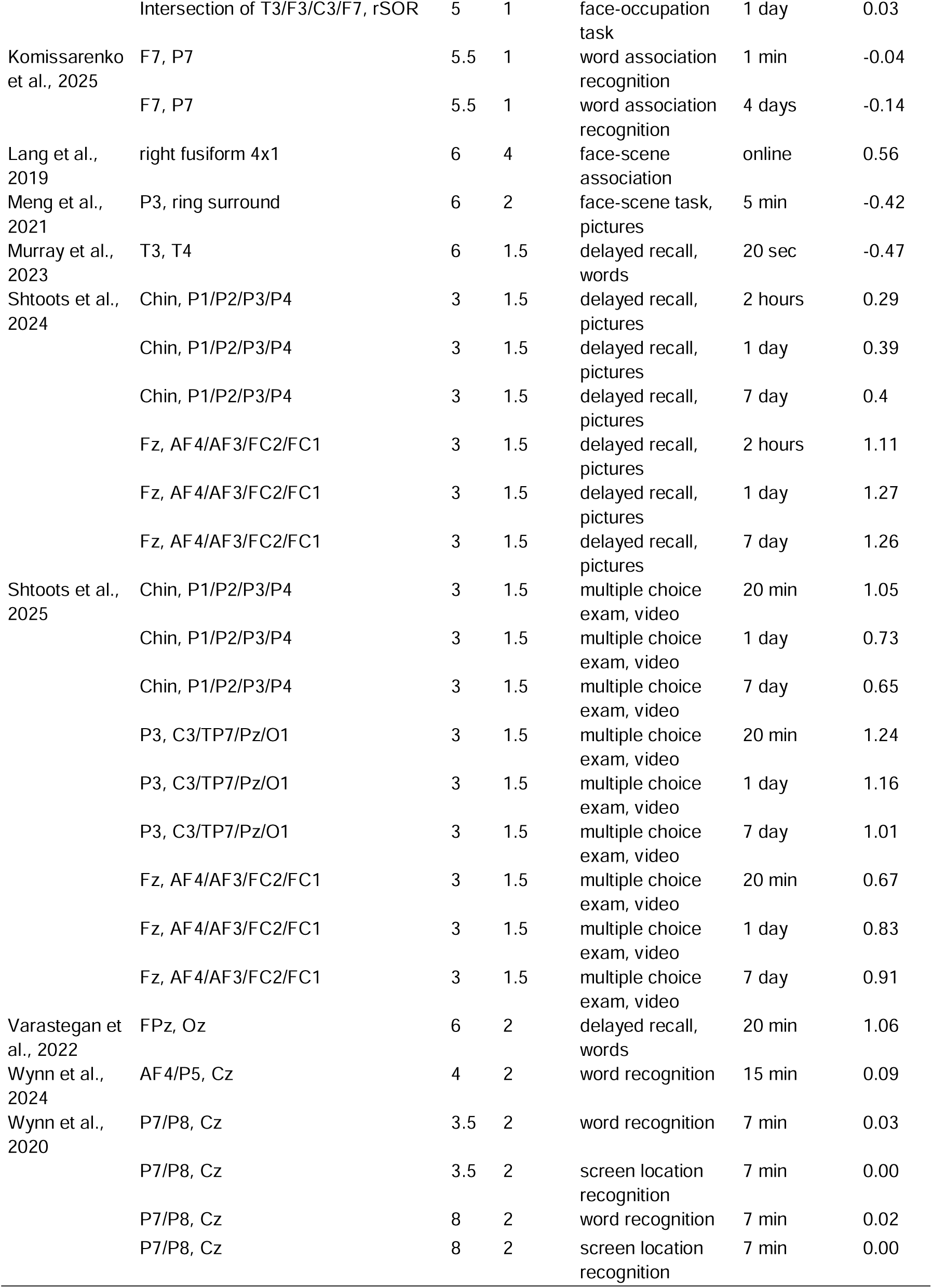
Overview of studies using theta tACS included in the meta-analysis.

**Table 2.**
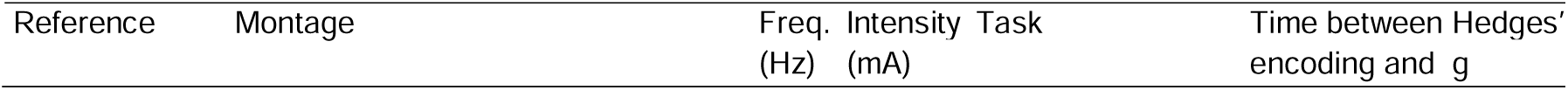
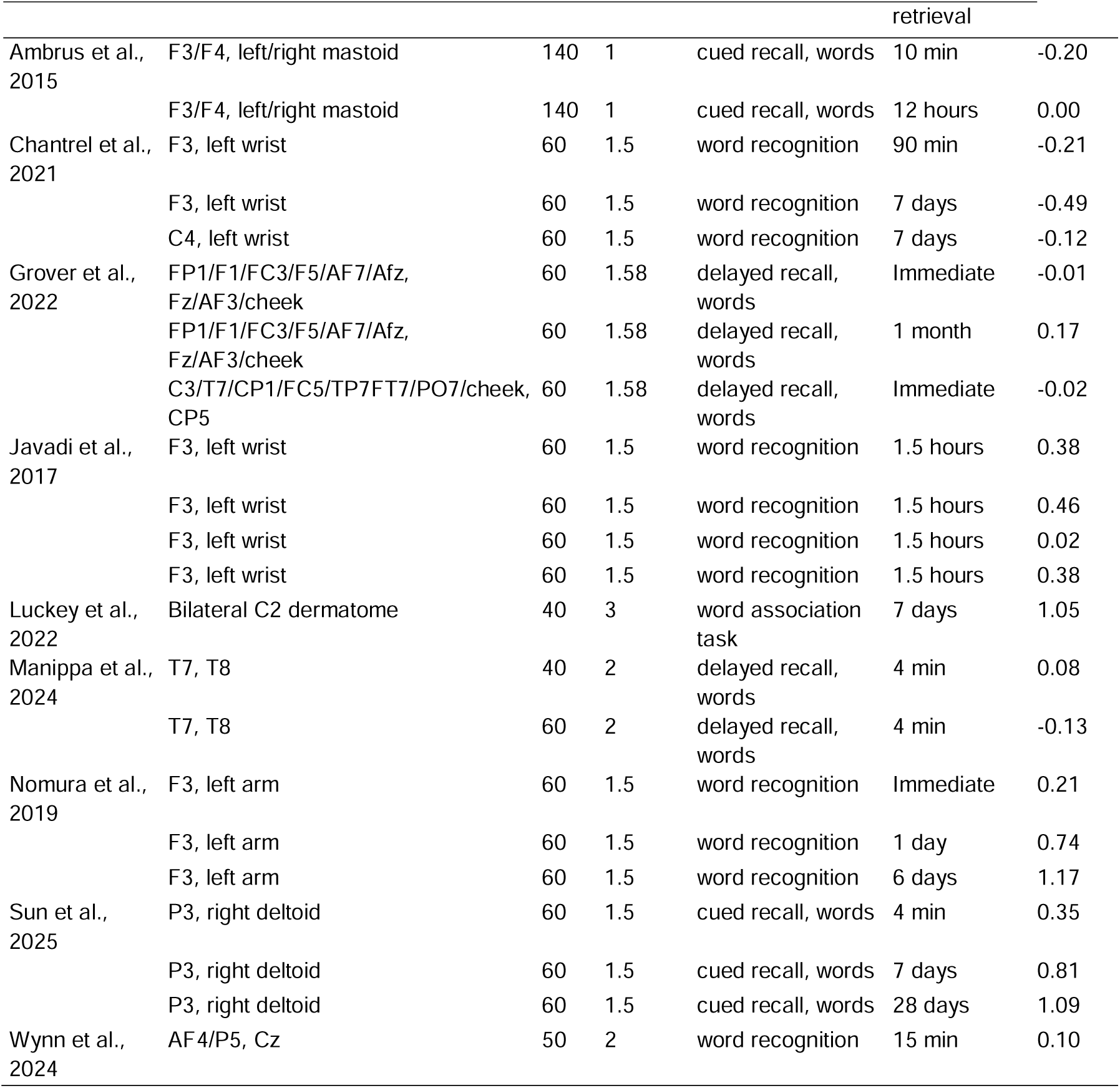
Overview of studies using gamma tACS included in the meta-analysis.

|  |  |  |  |  | retrieval |  |
| --- | --- | --- | --- | --- | --- | --- |
| Ambrus et al., 2015 | F3/F4, left/right mastoid | 140 | 1 | cued recall, words | 10 min | -0.20 |
|  | F3/F4, left/right mastoid | 140 | 1 | cued recall, words | 12 hours | 0.00 |
| Chantrel et al., 2021 | F3, left wrist | 60 | 1.5 | word recognition | 90 min | -0.21 |
|  | F3, left wrist | 60 | 1.5 | word recognition | 7 days | -0.49 |
|  | C4, left wrist | 60 | 1.5 | word recognition | 7 days | -0.12 |
| Grover et al., 2022 | FP1/F1/FC3/F5/AF7/Afz, Fz/AF3/cheek | 60 | 1.58 | delayed recall, words | Immediate | -0.01 |
|  | FP1/F1/FC3/F5/AF7/Afz, Fz/AF3/cheek | 60 | 1.58 | delayed recall, words | 1 month | 0.17 |
|  | C3/T7/CP1/FC5/TP7FT7/PO7/cheek, CP5 | 60 | 1.58 | delayed recall, words | Immediate | -0.02 |
| Javadi et al., 2017 | F3, left wrist | 60 | 1.5 | word recognition | 1.5 hours | 0.38 |
|  | F3, left wrist | 60 | 1.5 | word recognition | 1.5 hours | 0.46 |
|  | F3, left wrist | 60 | 1.5 | word recognition | 1.5 hours | 0.02 |
|  | F3, left wrist | 60 | 1.5 | word recognition | 1.5 hours | 0.38 |
| Luckey et al., 2022 | Bilateral C2 dermatome | 40 | 3 | word association task | 7 days | 1.05 |
| Manippa et al., 2024 | T7, T8 | 40 | 2 | delayed recall, words | 4 min | 0.08 |
|  | T7, T8 | 60 | 2 | delayed recall, words | 4 min | -0.13 |
| Nomura et al., 2019 | F3, left arm | 60 | 1.5 | word recognition | Immediate | 0.21 |
|  | F3, left arm | 60 | 1.5 | word recognition | 1 day | 0.74 |
|  | F3, left arm | 60 | 1.5 | word recognition | 6 days | 1.17 |
| Sun et al., 2025 | P3, right deltoid | 60 | 1.5 | cued recall, words | 4 min | 0.35 |
|  | P3, right deltoid | 60 | 1.5 | cued recall, words | 7 days | 0.81 |
|  | P3, right deltoid | 60 | 1.5 | cued recall, words | 28 days | 1.09 |
| Wynn et al., 2024 | AF4/P5, Cz | 50 | 2 | word recognition | 15 min | 0.10 |

## Results

In total 20 articles were identified for inclusion in the present meta-modeling analysis^25–28,33,56–70^. The analysis was split for studies that targeted theta oscillations (3-8 Hz) and gamma oscillations (30-200 Hz). For theta tACS studies, N=36, and for gamma tACS studies, N=22 effect sizes were obtained (Figure 2C). In three papers, tACS frequencies other than theta and gamma were studied, which were not included in our analysis (Supplementary Table 1). Since studies used various montages, we first aimed to get a sense of the general distribution of E-fields (Figure 2B). As such, we averaged E-fields across all studies (Figure 2A). For both frequencies, the simulations indicated widespread distribution across the brain, which is in line with electrodes being placed on frontal, temporal and occipito-parietal regions. However, for theta the averaged E-field was stronger at the left parietal cortex, whereas for gamma it was stronger for left frontal, temporal and parietal regions. This reflects a bias in the literature of preferentially targeting the left hemisphere.

**Figure 2.**
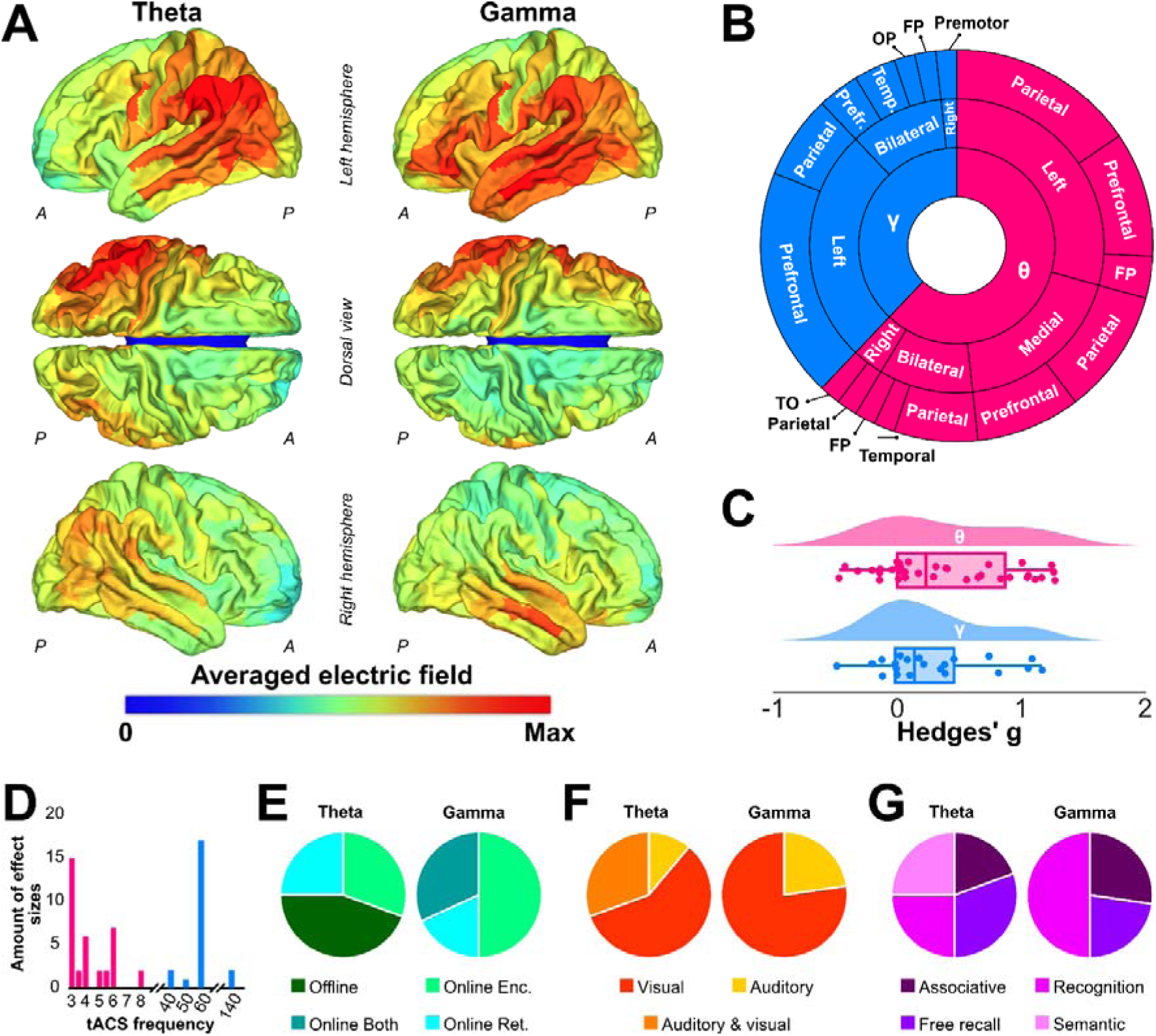
A) E-field strength averaged over all montages used in the selected articles, split for theta and gamma stimulation. Warm colors, where E-fields were strong on average, reflect regions commonly targeted across studies, while cold colors, where E-fields were weak on average, indicate regions less commonly targeted. B) A sunburst plot providing a global reflection of montages used across studies. FP: frontoparietal, OP: occipitoparietal, TO: Temporo-occipital. C) Raincloud distribution of effect sizes for theta and gamma tACS studies, showing an overall cumulative effect. Box plot limits correspond to the 25th and 75th percentiles, with the center line representing the median, and whiskers representing 1.5x interquartile range. D) The distribution of tACS frequencies used in the included studies. E) Pie charts showing whether studies applied tACS during encoding, retrieval, both, or offline (between encoding and retrieval). F) Pie charts showing how stimuli were presented across studies, either visually, auditorily, or a combination of both. G) Pie charts showing the type of LTM that was investigated across studies.

Across all included studies, a wide variety of study designs and tasks were used (Figure 2D-G). The majority of theta tACS studies used stimulation frequencies at the lower end of the theta range, with 75% of effect sizes from studies using a frequency ≤ 5.5 Hz. In those studies that applied gamma tACS, 60 Hz was the most common frequency (77.2%). For theta tACS studies, stimulation was either applied during encoding (30.6%), retrieval (25%), or offline (44.4%). For gamma tACS studies, stimulation was applied only during the task: During encoding (50%), retrieval (18.2%), or both (31.8%). Most studies used visual stimuli (theta: 58.3%; gamma: 77.2%), while the remaining studies used auditory stimuli (theta: 11.1%; gamma: 22.8%), or a combination of both (theta: 30.5%). Furthermore, various LTM task variations were used, including semantic memory, associative memory, free item recall, and item recognition (Figure 2G). Semantic memory studies provided to-be-memorized information that was subsequently quizzed. In studies using associative memory tasks, stimuli were paired, and the association between these items had to be remembered. In free recall tasks, the goal was to recall as many items from a given list as possible. In recognition tasks, a set of stimuli was memorized, and during retrieval, a set of new and old items was presented, with the goal of recognizing which items had been seen before. Given the wide variety of designs, it was not possible to include these factors in the subsequent meta-modeling analysis, as the small sample sizes would have yielded unreliable results.

### Theta tACS on LTM

For the theta tACS we found no regions where E-field strength showed a positive linear relationship with LTM performance, with a maximum of PEI = 0.052 (p = 0.761; right area 8B; frontal eye field). In contrast, E-field strength in the left frontal and temporal regions showed a negative linear association with LTM performance (p<0.001: N=2; 0.001<p<0.01: N=34; 0.01<p<0.05: N=39). Results are shown in Figure 3 and Supplementary Table 2. The strongest PEI values were found in regions 6r (PEI = -0.545, p <0.001) and 44 (PEI = -0.526, p < 0.001) of the HCP-MMP atlas, corresponding to the posterior part of the dorsolateral prefrontal and inferior frontal cortex, respectively.

**Figure 3.**
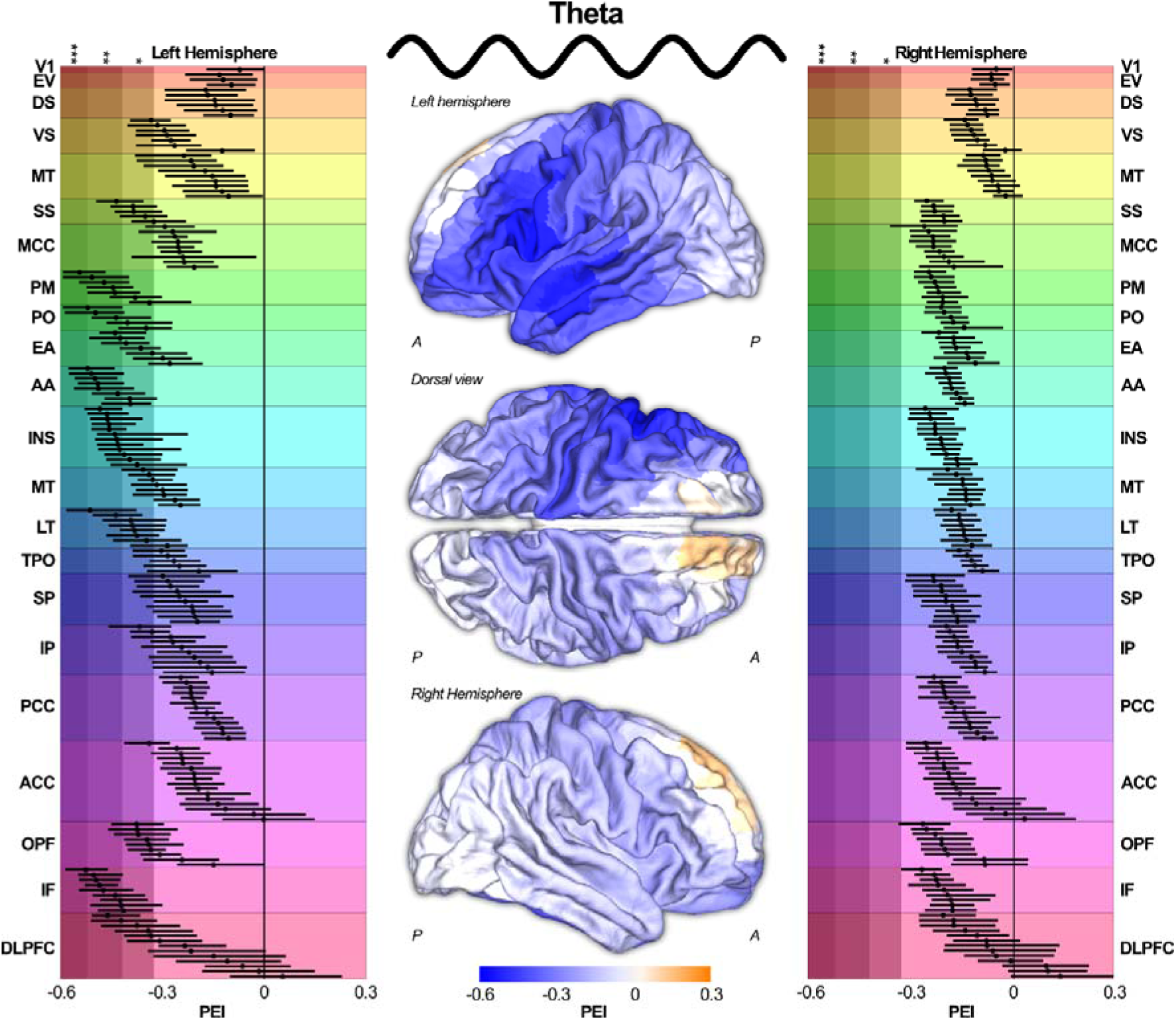
Distribution of PEI values across the brain related to theta tACS. Exact PEI values for all 360 regions, categorized into 22 larger regions (Huang et al., 2022; reflected by rainbow-colored shading), are reflected on the left and right. Gray shading reflects levels of significance with light shading p<0.05 (*), medium shading p<0.01 (**), and dark shading p<0.001 (***).

### Gamma tACS on LTM

For gamma tACS there was tentative evidence for E-field strength in bilateral occipital and parietal regions showing a positive linear relationship with LTM performance (p<0.05: N=24). The regions with the strongest associations between E-field strength and Hedges’ g were V1 (PEI = 0.514, p = 0.014) and V3A (PEI = 0.504, p = 0.016), in the visual cortex. No regions were found where E-field strength showed a significant negative linear relationship with performance, with a minimum of PEI = -0.232 (p = 0.297, left area anterior 9-46v; dorsolateral prefrontal cortex). Results are shown in Figure 4 and Supplementary Table 3.

**Figure 4.**
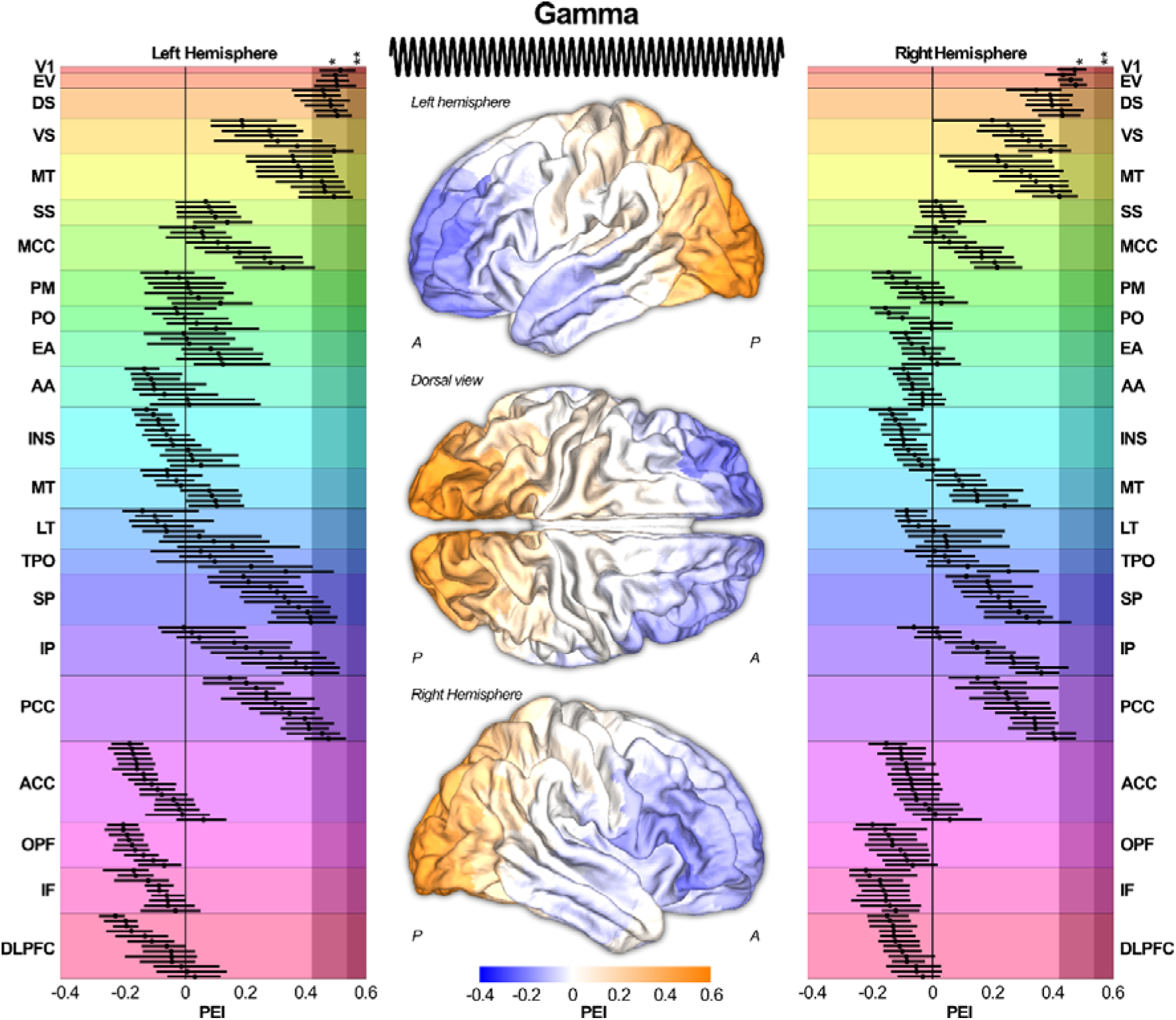
Distribution of PEI values across the brain related to gamma tACS. Exact PEI values for all 360 regions, categorized into 22 larger regions (Huang et al., 2022; reflected by rainbow-colored shading), are reflected on the left and right. Gray shading reflects levels of significance with light shading p<0.05 (*), medium shading p<0.01 (**), and dark shading p<0.001 (***).

### Quadratic associations between E-field magnitude and LTM

In an exploratory analysis we investigated whether regions show a quadratic relationship between E-field and effect size (Figure 5A). For theta tACS, we found that a quadratic curve explained the relationship between E-field strength and LTM performance better in 67 out of the 360 regions (i.e., a linear relationship explained more variance in the remaining 293 regions). This quadratic effect was observed in ventromedial prefrontal and medial temporal cortices. In all 67 regions a positive PEI was found ranging between PEI_QUAD_ = 0.349 (p = 0.039) and PEI_QUAD_ = 0.584 (p < 0.001). This positive relationship indicates that low and high E-field strength values had a positive effect on LTM performance, whereas medium E-field strength values did not. A visualization of this relationship is shown in Figure 5B, demonstrating the three regions with the highest PEI_QUAD_ values (Glasser region 47m, ventromedial prefrontal cortex, PEI_QUAD_ = 0.584; Glasser region PeEC, medial temporal cortex, PEI_QUAD_ = 0.547; and Glasser region PoI1, insular cortex, PEI_QUAD_ = 0.542). In these examples, it can be observed that at medium electric field strengths (≈0.05 to 0.1 mV/mm), tACS has no beneficial effect on LTM performance. However, at very small (<0.05 mV/mm) and large (>0.1 mV/mm) electric field strength values, there is. No significant negative quadratic PEI values were found. For gamma stimulation, no regions were identified where a quadratic fit was superior to a linear fit.

**Figure 5.**
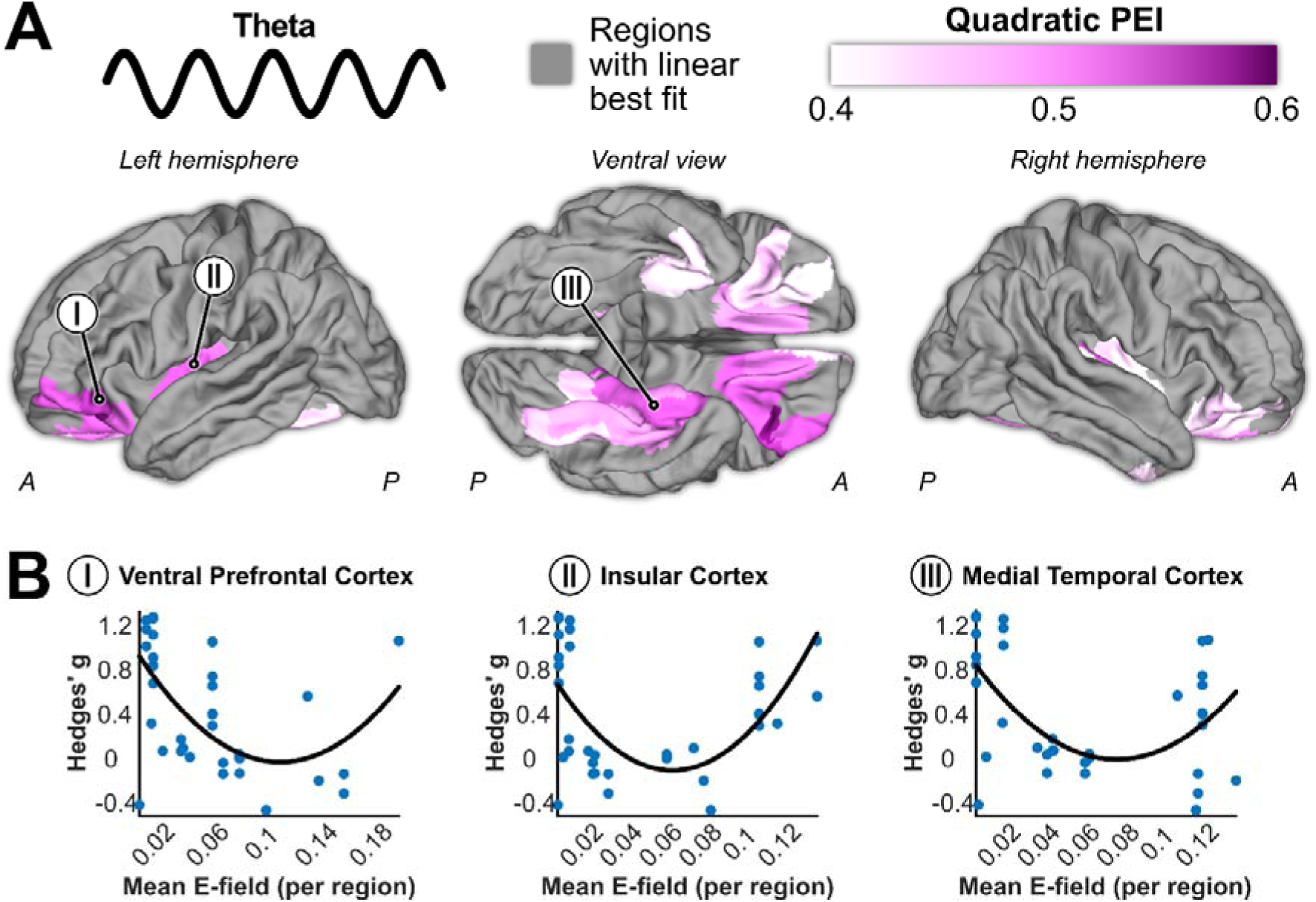
Distribution of PEI_QUAD_ values across the brain related to theta tACS, reflecting a quadratic relationship between E-field strength and behavioral performance.

## Discussion

We aimed to summarize published LTM-tACS studies to date by correlating tACS-induced electrical fields in brain models with corresponding changes in LTM performance. As a result, we found that within the theta frequency range, no regions showed a significant linear relationship that improved memory performance, but modulation of the left frontal and temporal lobes led to worse performance across studies. Exploring quadratic relationships, we found that theta over the ventromedial prefrontal and medial temporal cortices improves memory at low and high (but not moderate) E-field strength values. For gamma frequencies, parieto-occipital areas were associated with better performance, whereas frontal areas were associated with worse performance.

In contrast to our expectations, no clear linear positive relationship was found between theta stimulation and LTM performance. Only the dorsomedial prefrontal cortex showed a positive association. And while theta in the medial PFC has been linked to LTM in animal studies^12,71–73^, our results provide little support for this in humans. In contrast, and against our expectations, we found a negative association between LTM performance and E-field strength in left lateral prefrontal and temporal areas. The lateral prefrontal cortex is often associated with general cognitive performance and control, and various studies have at least suggested a potential role in LTM^25,74–77^. The temporal lobe includes and is connected to the hippocampus, which is often referred to as the main hub for LTM^78–81^. Within the hippocampus and medial temporal lobe, theta is a dominant oscillation^19,82–84^. Therefore, we hypothesized a positive PEI for this region. One possible explanation for this discrepancy is that the exogenous stimulation in the collected studies did not entrain but rather disrupted endogenous theta frequency or phase coupling. While tACS is generally thought to entrain regions and thereby strengthen endogenous oscillations^22^, various studies have emphasized that this may depend on specific circumstances^32^. For instance, several studies investigate in-phase vs anti-phase tACS, suggesting that in-phase stimulation is more likely to increase synchronization between regions^85–89^. Phase of the endogenous oscillations also matters. For instance, by using an amplitude-modulated tACS design and sophisticated artifact cleaning methods, Haslacher et al^90^ were able to record EEG while tACS is applied in a working memory task. They found that working memory depended on the phase lag between exogenous and endogenous theta oscillations. While this study looked at a different memory modality, it demonstrates the importance of phase alignment when aiming to modulate cognitive performance. Besides phase, recent studies have highlighted the importance of tACS intensity^52,53^. Specifically, lower intensities are more likely to decrease naturally occurring regional synchronization, rather than entrain a region. Furthermore, entrainment effects follow an Arnold’s tongue pattern, with the strongest entrainment occurring for stimulation frequencies that are close to endogenous rhythms^91,92^. This emphasizes the importance of personalized stimulation frequencies, which many of the here included studies did not apply^47^.

While we did not find any linear relationship between theta and LTM performance, when performing an additional analysis investigating a quadratic relationship, we found that this better explained the variance in 67 regions. Specifically, we found a significant positive quadratic effect in medial temporal and ventromedial prefrontal regions. In this case, a positive quadratic association reflects that the relation between E-field strength and LTM performance follows a U-shaped pattern. This suggests that parietal theta stimulation with low and high, but not moderate, E-field strength values was related to improved LTM performance. Indeed, various studies have demonstrated a nonlinear relationship between tACS intensity and neurophysiological and behavioral changes^51,54,55^. The U-shaped relationship between electric field strength and behavioral outcomes corresponds to physiological studies in animals showing that tACS-induced entrainment also follows a U-shape^52,93^. Krause et al^52^ found that while 2 mA tACS entrains neural firing to the exogenous current, 1 mA stimulation decreased entrainment compared to the natural entrainment to endogenous rhythms. This potentially suggests that tACS ‘makes things worse before it makes things better’. Nevertheless, the present results are surprising as they suggest improved LTM performance at electric field strength values < 0.05, which likely has a negligible effect on the brain^94^. As such, these effect sizes are more likely explained by factors unrelated to tACS, such as placebo-related or task-related learning effects. Alternatively, it could reflect a limitation of our method, as it does not take into account functional or structural connectivity between regions. For example, a region experiencing weak local electric fields may be strongly connected to another region exposed to higher field strengths, potentially leading to behavioral effects that the current method would not capture. Overall, while these interpretations remain speculative, the present findings highlight the importance of considering non-linear interactions between tACS and behavior..

For gamma tACS, we found that parieto-occipital regions were most strongly related to improvements in LTM. Indeed, studies have suggested that visual areas may serve as a storage or reactivation site for visual memory representations. For example, imaging work shows that visual memory traces may be maintained in the visual cortex^95–98^. Furthermore, gamma oscillations in particular have been related to sensory memory information^16,98,99^. On the other hand, gamma oscillations in the parieto-occipital cortex may reflect domain-general attentional mechanisms that support robust encoding and consolidation of memory by enhancing the binding of information and coordinating large-scale networks^100–106^. Previously, we found that gamma tACS of similar posterior regions enhances working memory performance^35^, suggesting that parieto-occipital gamma is not exclusively related to LTM. Rather, it may reflect a broader mechanism supporting cognitive operations across memory timescales^14^. In sensory cortices, gamma oscillations have been linked to attentional selection and feature binding, coordinating local neuronal ensembles to integrate information into coherent representations^101,107,108^. Thus, gamma tACS over parieto-occipital areas may facilitate long-term memory formation indirectly, by enhancing attentional and integrative processes in visual regions that support effective encoding and consolidation. Whether gamma tACS over the parietal cortex improves the sensory-mnemonic trace itself or improves the attentional processes cannot be deduced from our results. Nevertheless, our results suggest that stimulation of these regions most likely improves LTM performance.

When interpreting the main results, the variability in experimental designs was not considered a factor, as this would have been problematic with small sample sizes and would have led to unreliable results. Our PEI maps, therefore, reflect domain-general LTM. However, when interpreting our results, it is worth keeping these factors in mind. One such factor to consider is the exact frequency of stimulation. In working memory research, accumulating evidence suggests that stimulation at the lower range of theta may be beneficial for item maintenance^109–111^, although this observation likely also depends on the stimulation region^35^. Within the included studies, only Wynn et al^62^ compared low (3.5 Hz) and high (8 Hz) theta. In their visual recognition memory task, they found that neither stimulation frequency (applied over the parietal cortex) improved memory accuracy. However, low theta stimulation significantly reduced participants’ confidence in their memory performance. Evidence from intracranial recordings in surgical patients has shown that both low and high hippocampal theta power are positively related to spatial LTM performance^19^. Importantly, the majority of studies included in the present meta-analysis used low-frequency theta stimulation (<5.5 Hz). In how far this affected the results is open to debate, but it does reflect the need for more high-theta tACS studies and comparisons between low and high theta. In our meta-analysis, variation in the gamma frequency band was limited, with most studies using 60 Hz stimulation. While this reduces variability within our meta-analysis, it also highlights the importance of exploring a larger variety of stimulation frequencies within the gamma range.

The second factor to consider is the timing of stimulation. Across the studies, tACS was applied during encoding, during retrieval, during both encoding and retrieval, or offline (between encoding and retrieval). As such, the stimulation served conceptually distinct purposes, namely memory formation, consolidation, and recollection. The cortical networks and oscillatory processes related to these memory phases only partially overlap^112^, meaning that stimulation frequency and location need to be adjusted to each process. For instance, Shtoots and colleagues^27,68^ applied offline theta tACS (during the consolidation period) to the dorsomedial prefrontal cortex to test the role of midline theta in memory consolidation. On the other hand, Javadi et al^57^, who applied tACS during both encoding and retrieval, demonstrated the importance of matching stimulation frequencies for both periods, which is in line with the idea of context-dependent memory^113^. While speculative, it is possible that a specific memory phase is more susceptible to tACS than others. While our meta-analysis cannot answer this question, the different stimulation timings were represented by roughly equal numbers of effect sizes (Figure 2E), suggesting that our results are unlikely to be driven by a single factor.

A third factor that may influence the effect of tACS on LTM is the modality in which to-be-remembered stimuli are presented. Most studies used visual stimuli, with a minority of studies using auditory stimuli (or a combination of both). Other modalities, such as olfactory or tactile memory, were not investigated. This is a common bias for cognitive neuroscience in general, given the ease of computerized tasks^114^. We found gamma tACS over the occipitoparietal regions was positively correlated with improved LTM performance. A similar finding was previously observed for working memory^35^. More optimal visual processing, and thus increased occipital gamma, likely improves visual LTM performance. As such, it cannot be guaranteed that these results hold for other modalities.

A fourth factor to consider is the memory type. In other words, how participants had to remember information. Within our sample, some studies focused on the familiarity of items (e.g., “Have I seen this item before?”). Others targeted associative or semantic memory, requiring participants to retrieve relations between items or concepts embedded in a larger knowledge structure (e.g., “Did item X belong to item Y?”). This variety in forms of LTM is likely driven by different brain networks^112,115^. Furthermore, executive demands and cognitive control are likely to differ in associative memory tasks, compared to tasks with simple item recall or recognition^116^. Our results on theta tACS showed no significant regions positively associated with LTM performance. However, the regions showing a trend in this direction were in the prefrontal cortex. This pattern may be driven by studies employing tasks with higher cognitive demands, such as associative memory paradigms. Notably, the distribution of task types used in our sample was relatively balanced, making it unlikely that the overall results were dominated by a single form of memory. Nevertheless, future studies would benefit from directly comparing the effects of tACS across different forms of memory within the same experimental design.

Besides the above-mentioned factors some further considerations should be made when interpreting the results of the present study. First, given the reliance on existing data, not all potential montages have been explored. The PEI estimates rely on variability in E-fields and capture effects within these commonly stimulated areas, while regions rarely or never targeted should be interpreted with caution. The regions affected by the collected montages were rather evenly distributed across the brain. Nonetheless, there was some bias towards the temporoparietal areas and toward left hemispheric stimulation. Although such spatial bias could influence the results, the lack of overlap between the average E-field and PEI maps hints that any such effects were likely limited. Second, it is important to note that the results identify regions of interest based on the LTM-tACS studies conducted to date, not an exclusive map of regions essential for memory per se. Various regions might be crucial for LTM processing that are not susceptible to tACS-related improvements. Relatedly, since tACS targets primarily cortical regions, we can draw no conclusion about the causal influence of subcortical oscillations on LTM, for instance, those originating from hippocampal regions. Third, the present analyses were conducted using uncorrected p-values. This choice reflects a trade-off between statistical conservatism and sensitivity that is inherent to correlational meta-modeling approaches, which typically require large samples for stable inference. In the context of tACS research, the available literature remains limited in both the number of studies and outcome measures, constraining statistical power. Because the sample size in such analyses is determined by the existing literature rather than by an a priori design, conventional power calculations and standard multiple-comparison correction procedures are difficult to apply. While this approach increases sensitivity to detect potential effects, it also necessitates cautious interpretation of marginal findings. Fourth, electric field strength values were estimated using a set of 50 head models from the HCP S1200 dataset rather than participant-specific MRI data. This approach was chosen to incorporate realistic inter-individual anatomical variability in the absence of access to individual-level imaging data from the included studies. Nevertheless, this procedure constitutes an approximation, as the modeled electric fields cannot capture the exact anatomical properties of participants in the original experiments. Finally, as mentioned previously, our method does not consider the structural or functional connectivity and may be addressed by future refinements of electric field models In conclusion, we illustrate the potential of noninvasive brain stimulation to modulate LTM and shed light on the causal role of theta and gamma oscillations. As such, we find that tACS-related improvement to LTM is location and frequency-specific. Additionally, tACS-induced cognitive effects may not always scale linearly with intensity. Importantly, more studies are required to explore specific aspects of LTM, including different modalities and various methods of probing LTM. Nevertheless, we hope that these results provide a guide for future fundamental, translational, and therapeutic studies aimed at modulating memory.

## Conflicts of interest

The authors report no conflicts of interest.

## Funding

This work was supported by the XS grant of the Dutch Research Council (NWO; 406.XS.25.01.035).

## Author contributions

DC: Data curation, Formal analysis, Methodology, Writing - original draft. MW: Conceptualization, Formal analysis, Funding acquisition, Methodology, Supervision, Visualization, Writing - review & editing

## Notes

**Funding** This work was supported by the Dutch Research Council [406.XS.25.01.035].

**Conflicts of interest** The authors report no conflict of interests.

### Competing Interest Statement

The authors have declared no competing interest.

### Summary of Updates

To provide details on the literature search a PRISMA chart was added. Additionally, more information is added about the variability between study designs in stimulation frequency, online/offline stimulation, type of memory, and method of recollection. Finally, the quadratic analysis was modified and corrected.

## References

1. Squire LR, Wixted JT. The Cognitive Neuroscience of Human Memory Since H.M. Annual Review of Neuroscience. 2011;34(Volume 34, 2011):259-288. doi:10.1146/annurev-neuro-061010-113720

2. Jeneson A, Squire LR. Working memory, long-term memory, and medial temporal lobe function. Learn Mem. 2012;19(1):15–25. doi:10.1101/lm.024018.111

3. Pittenger C. Disorders of memory and plasticity in psychiatric disease. Dialogues in Clinical Neuroscience. 2013;15(4):455–463. doi:10.31887/DCNS.2013.15.4/cpittenger

4. Yang S, Yi YG, Chang MC. The effect of transcranial alternating current stimulation on functional recovery in patients with stroke: a narrative review. Front Neurol. 2023;14:1327383. doi:10.3389/fneur.2023.1327383

5. Daume J, Kamiński J, Schjetnan AGP, et al. Control of working memory by phase–amplitude coupling of human hippocampal neurons. Nature. 2024;629(8011):393–401. doi:10.1038/s41586-024-07309-z

6. Sawangjit A, Oyanedel CN, Niethard N, Salazar C, Born J, Inostroza M. The hippocampus is crucial for forming non-hippocampal long-term memory during sleep. Nature. 2018;564(7734):109–113. doi:10.1038/s41586-018-0716-8

7. Jetter W, Poser U, Freeman RB, Markowitsch HJ. A Verbal Long Term Memory Deficit in Frontal Lobe Damaged Patients. Cortex. 1986;22(2):229–242. doi:10.1016/S0010-9452(86)80047-8

8. Kesner RP. The posterior parietal cortex and long-term memory representation of spatial information. Neurobiology of Learning and Memory. 2009;91(2):197–206. doi:10.1016/j.nlm.2008.09.004

9. Olson IR, Berryhill M. Some surprising findings on the involvement of the parietal lobe in human memory. Neurobiology of Learning and Memory. 2009;91(2):155–165. doi:10.1016/j.nlm.2008.09.006

10. Squire LR. Memory for relations in the short term and the long term after medial temporal lobe damage. Hippocampus. 2017;27(5):608–612. doi:10.1002/hipo.22716

11. Miller JA, Tambini A, Kiyonaga A, D’Esposito M. Long-term learning transforms prefrontal cortex representations during working memory. Neuron. 2022;110(22):3805–3819.e6. doi:10.1016/j.neuron.2022.09.019

12. Sheynikhovich D, Otani S, Bai J, Arleo A. Long-term memory, synaptic plasticity and dopamine in rodent medial prefrontal cortex: Role in executive functions. Front Behav Neurosci. 2023;16. doi:10.3389/fnbeh.2022.1068271

13. Axmacher N, Mormann F, Fernández G, Elger CE, Fell J. Memory formation by neuronal synchronization. Brain Res Rev. 2006;52(1):170–182. doi:10.1016/j.brainresrev.2006.01.007

14. Nyhus E, Curran T. Functional Role of Gamma and Theta Oscillations in Episodic Memory. Neurosci Biobehav Rev. 2010;34(7):1023–1035. doi:10.1016/j.neubiorev.2009.12.014

15. Köster M. The theta-gamma code in predictive processing and mnemonic updating. Neuroscience & Biobehavioral Reviews. 2024;158:105529. doi:10.1016/j.neubiorev.2023.105529

16. Osipova D, Takashima A, Oostenveld R, Fernández G, Maris E, Jensen O. Theta and Gamma Oscillations Predict Encoding and Retrieval of Declarative Memory. J Neurosci. 2006;26(28):7523–7531. doi:10.1523/JNEUROSCI.1948-06.2006

17. Sederberg PB, Kahana MJ, Howard MW, Donner EJ, Madsen JR. Theta and Gamma Oscillations during Encoding Predict Subsequent Recall. J Neurosci. 2003;23(34):10809–10814. doi:10.1523/JNEUROSCI.23-34-10809.2003

18. Sederberg PB, Schulze-Bonhage A, Madsen JR, et al. Hippocampal and Neocortical Gamma Oscillations Predict Memory Formation in Humans. Cereb Cortex. 2007;17(5):1190–1196. doi:10.1093/cercor/bhl030

19. Vivekananda U, Bush D, Bisby JA, et al. Theta power and theta-gamma coupling support long-term spatial memory retrieval. Hippocampus. 2021;31(2):213–220. doi:10.1002/hipo.23284

20. Hanouneh S, Amin HU, Saad NM, Malik AS. EEG Power and Functional Connectivity Correlates with Semantic Long-Term Memory Retrieval. IEEE Access. 2018;6:8695–8703. doi:10.1109/ACCESS.2017.2788859

21. Scholz S, Schneider SL, Rose M. Differential effects of ongoing EEG beta and theta power on memory formation. PLOS ONE. 2017;12(2):e0171913. doi:10.1371/journal.pone.0171913

22. Polanía R, Nitsche MA, Ruff CC. Studying and modifying brain function with non-invasive brain stimulation. Nat Neurosci. 2018;21(2):174–187. doi:10.1038/s41593-017-0054-4

23. Elyamany O, Leicht G, Herrmann CS, Mulert C. Transcranial alternating current stimulation (tACS): from basic mechanisms towards first applications in psychiatry. Eur Arch Psychiatry Clin Neurosci. 2021;271(1):135–156. doi:10.1007/s00406-020-01209-9

24. Agboada D, Zhao Z, Wischnewski M. Neuroplastic effects of transcranial alternating current stimulation (tACS): from mechanisms to clinical trials. Front Hum Neurosci. 2025;19. doi:10.3389/fnhum.2025.1548478

25. Grover S, Wen W, Viswanathan V, Gill CT, Reinhart RMG. Long-lasting, dissociable improvements in working memory and long-term memory in older adults with repetitive neuromodulation. Nat Neurosci. 2022;25:1237–1246. doi:10.1038/s41593-022-01132-3

26. Manippa V, Nitsche MA, Filardi M, et al. Temporal gamma tACS and auditory stimulation affect verbal memory in healthy adults. Psychophysiology. 2024;61(11):e14653. doi:10.1111/psyp.14653

27. Shtoots L, Nadler A, Partouche R, et al. Frontal midline theta transcranial alternating current stimulation enhances early consolidation of episodic memory. npj Sci Learn. 2024;9(1):8. doi:10.1038/s41539-024-00222-0

28. Wynn SC, Townsend CD, Nyhus E. The role of theta and gamma oscillations in item memory, source memory, and memory confidence. Psychophysiology. 2024;61(9):e14602. doi:10.1111/psyp.14602

29. Krause MR, Vieira PG, Csorba BA, Pilly PK, Pack CC. Transcranial alternating current stimulation entrains single-neuron activity in the primate brain. Proc Natl Acad Sci U S A. 2019;116(12):5747–5755. doi:10.1073/pnas.1815958116

30. Wischnewski M, Tran H, Zhao Z, et al. Induced neural phase precession through exogenous electric fields. Nat Commun. 2024;15(1):1687. doi:10.1038/s41467-024-45898-5

31. Schutter DJLG, Wischnewski M. A meta-analytic study of exogenous oscillatory electric potentials in neuroenhancement. Neuropsychologia. 2016;86:110–118. doi:10.1016/j.neuropsychologia.2016.04.011

32. Wischnewski M, Alekseichuk I, Opitz A. Neurocognitive, physiological, and biophysical effects of transcranial alternating current stimulation. Trends in Cognitive Sciences. 2023;27(2):189–205. doi:10.1016/j.tics.2022.11.013

33. Alekseichuk I, Turi Z, Veit S, Paulus W. Model-driven neuromodulation of the right posterior region promotes encoding of long-term memories. Brain Stimulation. 2020;13(2):474–483. doi:10.1016/j.brs.2019.12.019

34. Wischnewski M, Mantell KE, Opitz A. Identifying regions in prefrontal cortex related to working memory improvement: A novel meta-analytic method using electric field modeling. Neuroscience & Biobehavioral Reviews. 2021;130:147–161.

35. Wischnewski M, Berger TA, Opitz A, Alekseichuk I. Causal functional maps of brain rhythms in working memory. Proc Natl Acad Sci USA. 2024;121(14):e2318528121. doi:10.1073/pnas.2318528121

36. Wischnewski M, Berger TA, Opitz A. Meta-modeling the effects of anodal left prefrontal transcranial direct current stimulation on working memory performance. Imaging Neuroscience. 2024;2:1–14. doi:10.1162/imag_a_00078

37. Yachou Y, Bouaziz N, Makdah G, et al. Transcranial direct current stimulation in patients with depression: An electric field modeling meta-analysis. Journal of Affective Disorders. 2025;374:540–552. doi:10.1016/j.jad.2025.01.001

38. Thielscher A, Antunes A, Saturnino GB. Field modeling for transcranial magnetic stimulation: A useful tool to understand the physiological effects of TMS? In: 2015 37th Annual International Conference of the IEEE Engineering in Medicine and Biology Society (EMBC). 2015:222-225. doi:10.1109/EMBC.2015.7318340

39. Moher D, Liberati A, Tetzlaff J, Altman DG. Preferred reporting items for systematic reviews and meta-analyses: the PRISMA statement. BMJ. 2009;339:b2535. doi:10.1136/bmj.b2535

40. Hedges LV, Olkin I. Statistical Methods for Meta-Analysis. Academic Press; 1985.

41. Cohen J. Statistical Power Analysis for the Behavioral Sciences. Academic Press; 1977.

42. Van Essen DC, Smith SM, Barch DM, et al. The WU-Minn human connectome project: an overview. Neuroimage. 2013;80:62–79. Accessed October 24, 2025. https://www.sciencedirect.com/science/article/pii/S1053811913005351

43. Berger TA, Wischnewski M, Opitz A, Alekseichuk I. Human head models and populational framework for simulating brain stimulations. Sci Data. 2025;12(1):516. doi:10.1038/s41597-025-04886-0

44. Windhoff M, Opitz A, Thielscher A. Electric field calculations in brain stimulation based on finite elements: an optimized processing pipeline for the generation and usage of accurate individual head models. Hum Brain Mapp. 2013;34(4):923–935. doi:10.1002/hbm.21479

45. McCann H, Pisano G, Beltrachini L. Variation in reported human head tissue electrical conductivity values. Brain Topogr. 2019;32(5):825–858. doi:10.1007/s10548-019-00710-2

46. Katoch N, Kim Y, Choi BK, et al. Estimation of brain tissue response by electrical stimulation in a subject-specific model implemented by conductivity tensor imaging. Front Neurosci. 2023;17. doi:10.3389/fnins.2023.1197452

47. Hoornweder SV, Stagg CJ, Wischnewski M. Personalizing transcranial electrical stimulation. Trends in Neurosciences. 2025;0(0). doi:10.1016/j.tins.2025.07.007

48. Glasser MF, Coalson TS, Robinson EC, et al. A multi-modal parcellation of human cerebral cortex. Nature. 2016;536(7615):171–178. doi:10.1038/nature18933

49. Glasser MF, Smith SM, Marcus DS, et al. The Human Connectome Project’s neuroimaging approach. Nat Neurosci. 2016;19(9):1175–1187. doi:10.1038/nn.4361

50. Huang CC, Rolls ET, Feng J, Lin CP. An extended Human Connectome Project multimodal parcellation atlas of the human cortex and subcortical areas. Brain Struct Funct. 2022;227(3):763–778. doi:10.1007/s00429-021-02421-6

51. Moliadze V, Atalay D, Antal A, Paulus W. Close to threshold transcranial electrical stimulation preferentially activates inhibitory networks before switching to excitation with higher intensities. Brain Stimulation. 2012;5(4):505–511. doi:10.1016/j.brs.2011.11.004

52. Krause MR, Vieira PG, Thivierge JP, Pack CC. Brain stimulation competes with ongoing oscillations for control of spike timing in the primate brain. PLOS Biology. 2022;20(5):e3001650. doi:10.1371/journal.pbio.3001650

53. Zhao Z, Shirinpour S, Tran H, Wischnewski M, Opitz A. Intensity- and frequency-specific effects of transcranial alternating current stimulation are explained by network dynamics. J Neural Eng. 2024;21(2):026024. doi:10.1088/1741-2552/ad37d9

54. Hsu WY, Zanto T, Park JE, Gazzaley A, Bove RM. Effects of transcranial alternating current stimulation on cognitive function in people with multiple sclerosis: A randomized controlled trial. Mult Scler Relat Disord. 2023;80:105090. doi:10.1016/j.msard.2023.105090

55. Wansbrough K, Marinovic W, Fujiyama H, Vallence AM. Beta tACS of varying intensities differentially affect resting-state and movement-related M1-M1 connectivity. Front Neurosci. 2024;18. doi:10.3389/fnins.2024.1425527

56. Ambrus GG, Pisoni A, Primaßin A, Turi Z, Paulus W, Antal A. Bi-frontal transcranial alternating current stimulation in the ripple range reduced overnight forgetting. Front Cell Neurosci. 2015;9. doi:10.3389/fncel.2015.00374

57. Javadi AH, Glen JC, Halkiopoulos S, Schulz M, Spiers HJ. Oscillatory Reinstatement Enhances Declarative Memory. J Neurosci. 2017;37(41):9939–9944. doi:10.1523/JNEUROSCI.0265-17.2017

58. Lang S, Gan LS, Alrazi T, Monchi O. Theta band high definition transcranial alternating current stimulation, but not transcranial direct current stimulation, improves associative memory performance. Sci Rep. 2019;9(1):8562. doi:10.1038/s41598-019-44680-8

59. Nomura T, Asao A, Kumasaka A. Transcranial alternating current stimulation over the prefrontal cortex enhances episodic memory recognition. Exp Brain Res. 2019;237(7):1709–1715. doi:10.1007/s00221-019-05543-w

60. Ergo K, Loof ED, Debra G, Pastötter B, Verguts T. Failure to modulate reward prediction errors in declarative learning with theta (6 Hz) frequency transcranial alternating current stimulation. PLOS ONE. 2020;15(12):e0237829. doi:10.1371/journal.pone.0237829

61. Klink K, Peter J, Wyss P, Klöppel S. Transcranial Electric Current Stimulation During Associative Memory Encoding: Comparing tACS and tDCS Effects in Healthy Aging. Front Aging Neurosci. 2020;12. doi:10.3389/fnagi.2020.00066

62. Wynn SC, Kessels RPC, Schutter DJLG. Effects of parietal exogenous oscillatory field potentials on subjectively perceived memory confidence. Neurobiology of Learning and Memory. 2020;168:107140. doi:10.1016/j.nlm.2019.107140

63. Chantrel Y, Trabattoni V, Orton L, Javadi AH. Intrinsic reinstatement of induced oscillatory context. bioRxiv. Preprint posted online January 26, 2021:2021.01.25.428096. doi:10.1101/2021.01.25.428096

64. Meng A, Kaiser M, de Graaf TA, et al. Transcranial alternating current stimulation at theta frequency to left parietal cortex impairs associative, but not perceptual, memory encoding. Neurobiology of Learning and Memory. 2021;182:107444. doi:10.1016/j.nlm.2021.107444

65. Luckey AM, McLeod SL, Mohan A, Vanneste S. Potential role for peripheral nerve stimulation on learning and long-term memory: A comparison of alternating and direct current stimulations. Brain Stimulation: Basic, Translational, and Clinical Research in Neuromodulation. 2022;15(3):536–545. doi:10.1016/j.brs.2022.03.001

66. Murray NWG, Graham PL, Sowman PF, Savage G. Theta tACS impairs episodic memory more than tDCS. Sci Rep. 2023;13(1):716. doi:10.1038/s41598-022-27190-y

67. Varastegan S, Kazemi R, Rostami R, Khomami S, Zandbagleh A, Hadipour AL. Remember NIBS? tACS improves memory performance in elders with subjective memory complaints. GeroScience. 2023;45(2):851–869. doi:10.1007/s11357-022-00677-2

68. Shtoots L, Barzilay R, Gigi T, Kostovetsky V, Pollock A, Levy DA. Theta Stimulation Enhances Early Consolidation of Semantic Memory. J Cogn Neurosci. 2025;37(9):1496–1510. doi:10.1162/jocn_a_02322

69. Komissarenko A, Stupina E, Malyutina S. Targeting the neural bases of novel word acquisition using theta-band transcranial alternating current stimulation. Language, Cognition and Neuroscience. 2025;40(4):487–507. doi:10.1080/23273798.2025.2454001

70. Sun S, Annaka H, Nomura T. Gamma-frequency transcranial alternating current stimulation over the left posterior parietal cortex enhances the long-term retention of associative memory. Exp Brain Res. 2025;243(3):62. doi:10.1007/s00221-025-07009-8

71. Miller EK, Cohen JD. An integrative theory of prefrontal cortex function. Annu Rev Neurosci. 2001;24:167–202. doi:10.1146/annurev.neuro.24.1.167

72. Euston DR, Gruber AJ, McNaughton BL. The Role of Medial Prefrontal Cortex in Memory and Decision Making. Neuron. 2012;76(6):1057–1070. doi:10.1016/j.neuron.2012.12.002

73. Martínez MC, Villar ME, Ballarini F, Viola H. Retroactive interference of object-in-context long-term memory: Role of dorsal hippocampus and medial prefrontal cortex. Hippocampus. 2014;24(12):1482–1492. doi:10.1002/hipo.22328

74. Blumenfeld RS, Lee TG, D’Esposito M. The effects of lateral prefrontal transcranial magnetic stimulation on item memory encoding. Neuropsychologia. 2014;53:197–202. doi:10.1016/j.neuropsychologia.2013.11.021

75. Ghazizadeh A, Hong S, Hikosaka O. Prefrontal Cortex Represents Long-Term Memory of Object Values for Months. Current Biology. 2018;28(14):2206–2217.e5. doi:10.1016/j.cub.2018.05.017

76. Ghazizadeh A, Griggs W, Leopold DA, Hikosaka O. Temporal–prefrontal cortical network for discrimination of valuable objects in long-term memory. Proceedings of the National Academy of Sciences. 2018;115(9):E2135–E2144. doi:10.1073/pnas.1707695115

77. Blumenfeld RS, Ranganath C. Chapter 12 - The lateral prefrontal cortex and human long-term memory. In: D’Esposito M, Grafman JH, eds. Handbook of Clinical Neurology. Vol 163. The Frontal Lobes. Elsevier; 2019:221-235. doi:10.1016/B978-0-12-804281-6.00012-4

78. Treves A, Rolls ET. Computational analysis of the role of the hippocampus in memory. Hippocampus. 1994;4(3):374–391. doi:10.1002/hipo.450040319

79. Simons JS, Spiers HJ. Prefrontal and medial temporal lobe interactions in long-term memory. Nat Rev Neurosci. 2003;4(8):637–648. doi:10.1038/nrn1178

80. Squire LR, Stark CEL, Clark RE. The medial temporal lobe. Annual Review of Neuroscience. 2004;27(Volume 27, 2004):279-306. doi:10.1146/annurev.neuro.27.070203.144130

81. Lisman JE, Grace AA. The Hippocampal-VTA Loop: Controlling the Entry of Information into Long-Term Memory. Neuron. 2005;46(5):703–713. doi:10.1016/j.neuron.2005.05.002

82. Guderian S, Schott BH, Richardson-Klavehn A, Düzel E. Medial temporal theta state before an event predicts episodic encoding success in humans. Proceedings of the National Academy of Sciences. 2009;106(13):5365–5370. doi:10.1073/pnas.0900289106

83. Lega BC, Jacobs J, Kahana M. Human hippocampal theta oscillations and the formation of episodic memories. Hippocampus. 2012;22(4):748–761. doi:10.1002/hipo.20937

84. Solomon EA, Stein JM, Das S, et al. Dynamic Theta Networks in the Human Medial Temporal Lobe Support Episodic Memory. Current Biology. 2019;29(7):1100–1111.e4. doi:10.1016/j.cub.2019.02.020

85. Polanía R, Nitsche MA, Korman C, Batsikadze G, Paulus W. The importance of timing in segregated theta phase-coupling for cognitive performance. Current Biology. 2012;22(14):1314–1318. doi:10.1016/j.cub.2012.05.021

86. Violante IR, Li LM, Carmichael DW, et al. Externally induced frontoparietal synchronization modulates network dynamics and enhances working memory performance. eLife. 2017;6:e22001. doi:10.7554/eLife.22001

87. Schwab BC, Misselhorn J, Engel AK. Modulation of large-scale cortical coupling by transcranial alternating current stimulation. Brain Stimulation. 2019;12(5):1187–1196. doi:10.1016/j.brs.2019.04.013

88. Schwab BC, König P, Engel AK. Spike-timing-dependent plasticity can account for connectivity aftereffects of dual-site transcranial alternating current stimulation. NeuroImage. 2021;237:118179. doi:10.1016/j.neuroimage.2021.118179

89. Barzegar S, Kakies CFM, Ciuperc= D, Wischnewski M. Transcranial alternating current stimulation for investigating complex oscillatory dynamics and interactions. International Journal of Psychophysiology. 2025;212:112579. doi:10.1016/j.ijpsycho.2025.112579

90. Haslacher D, Cavallo A, Reber P, et al. Working memory enhancement using real-time phase-tuned transcranial alternating current stimulation. Brain Stimulation. 2024;17(4):850–859. doi:10.1016/j.brs.2024.07.007

91. Herrmann CS, Rach S, Neuling T, Strüber D. Transcranial alternating current stimulation: a review of the underlying mechanisms and modulation of cognitive processes. Frontiers in human neuroscience. 2013;7:279. Accessed October 15, 2025. https://www.frontiersin.org/journals/human-neuroscience/articles/10.3389/fnhum.2013.00279/full

92. Antal A, Herrmann CS. Transcranial Alternating Current and Random Noise Stimulation: Possible Mechanisms. Neural Plasticity. 2016;2016(1):3616807. doi:10.1155/2016/3616807

93. Vieira PG, Krause MR, Pack CC. Temporal interference stimulation disrupts spike timing in the primate brain. Nat Commun. 2024;15(1):4558. doi:10.1038/s41467-024-48962-2

94. Alekseichuk I, Wischnewski M, Opitz A. A minimum effective dose for (transcranial) alternating current stimulation. Brain Stimulation: Basic, Translational, and Clinical Research in Neuromodulation. 2022;15(5):1221–1222. doi:10.1016/j.brs.2022.08.018

95. Wheeler ME, Petersen SE, Buckner RL. Memory’s echo: Vivid remembering reactivates sensory-specific cortex. Proceedings of the National Academy of Sciences. 2000;97(20):11125–11129. doi:10.1073/pnas.97.20.11125

96. Fedurco M. Long-Term Memory Search across the Visual Brain. Neural Plast. 2012;2012:392695. doi:10.1155/2012/392695

97. Favila SE, Kuhl BA, Winawer J. Perception and memory have distinct spatial tuning properties in human visual cortex. Nat Commun. 2022;13(1):5864. doi:10.1038/s41467-022-33161-8

98. Griffiths BJ, Jensen O. Gamma oscillations and episodic memory. Trends in Neurosciences. 2023;46(10):832-846. doi:10.1016/j.tins.2023.07.003

99. Köster M, Gruber T. Rhythms of human attention and memory: An embedded process perspective. Front Hum Neurosci. 2022;16. doi:10.3389/fnhum.2022.905837

100. Keizer AW, Verment RS, Hommel B. Enhancing cognitive control through neurofeedback: A role of gamma-band activity in managing episodic retrieval. NeuroImage. 2010;49(4):3404–3413. doi:10.1016/j.neuroimage.2009.11.023

101. Morgan HM, Muthukumaraswamy SD, Hibbs CS, et al. Feature integration in visual working memory: parietal gamma activity is related to cognitive coordination. Journal of Neurophysiology. 2011;106(6):3185–3194. doi:10.1152/jn.00246.2011

102. Rouhinen S, Panula J, Palva JM, Palva S. Load Dependence of β and γ Oscillations Predicts Individual Capacity of Visual Attention. J Neurosci. 2013;33(48):19023–19033. doi:10.1523/JNEUROSCI.1666-13.2013

103. Tseng P, Chang YT, Chang CF, Liang WK, Juan CH. The critical role of phase difference in gamma oscillation within the temporoparietal network for binding visual working memory. Sci Rep. 2016;6(1):32138. doi:10.1038/srep32138

104. Magazzini L, Singh KD. Spatial attention modulates visual gamma oscillations across the human ventral stream. NeuroImage. 2018;166:219–229. doi:10.1016/j.neuroimage.2017.10.069

105. Marshall TR, den Boer S, Cools R, Jensen O, Fallon SJ, Zumer JM. Occipital Alpha and Gamma Oscillations Support Complementary Mechanisms for Processing Stimulus Value Associations. J Cogn Neurosci. 2018;30(1):119–129. doi:10.1162/jocn_a_01185

106. Kasten FH, Wendeln T, Stecher HI, Herrmann CS. Hemisphere-specific, differential effects of lateralized, occipital–parietal α- versus γ-tACS on endogenous but not exogenous visual-spatial attention. Sci Rep. 2020;10(1):12270. doi:10.1038/s41598-020-68992-2

107. Tiesinga P, Sejnowski TJ. Cortical Enlightenment: Are Attentional Gamma Oscillations Driven by ING or PING? Neuron. 2009;63(6):727–732. doi:10.1016/j.neuron.2009.09.009

108. Misselhorn J, Schwab BC, Schneider TR, Engel AK. Synchronization of Sensory Gamma Oscillations Promotes Multisensory Communication. eNeuro. 2019;6(5). doi:10.1523/ENEURO.0101-19.2019

109. Aktürk T, de Graaf TA, Güntekin B, Hanoğlu L, Sack AT. Enhancing memory capacity by experimentally slowing theta frequency oscillations using combined EEG-tACS. Sci Rep. 2022;12(1):14199. doi:10.1038/s41598-022-18665-z

110. Wolinski N, Cooper NR, Sauseng P, Romei V. The speed of parietal theta frequency drives visuospatial working memory capacity. PLOS Biology. 2018;16(3):e2005348. doi:10.1371/journal.pbio.2005348

111. Bender M, Romei V, Sauseng P. Slow Theta tACS of the Right Parietal Cortex Enhances Contralateral Visual Working Memory Capacity. Brain Topogr. 2019;32(3):477–481. doi:10.1007/s10548-019-00702-2

112. Ranganath C, Ritchey M. Two cortical systems for memory-guided behaviour. Nat Rev Neurosci. 2012;13(10):713–726. doi:10.1038/nrn3338

113. Heald JB, Lengyel M, Wolpert DM. Contextual inference in learning and memory. Trends in Cognitive Sciences. 2023;27(1):43–64. doi:10.1016/j.tics.2022.10.004

114. Hutmacher F. Why Is There So Much More Research on Vision Than on Any Other Sensory Modality? Front Psychol. 2019;10:2246. doi:10.3389/fpsyg.2019.02246

115. Eichenbaum H, Yonelinas AP, Ranganath C. The Medial Temporal Lobe and Recognition Memory. Annual Review of Neuroscience. 2007;30(Volume 30, 2007):123-152. doi:10.1146/annurev.neuro.30.051606.094328

116. Blumenfeld RS, Ranganath C. Prefrontal Cortex and Long-Term Memory Encoding: An Integrative Review of Findings from Neuropsychology and Neuroimaging. Neuroscientist. 2007;13(3):280–291. doi:10.1177/1073858407299290

