## Supplementary for "Mapping long-term memory modulation with transcranial alternating current stimulation"

| Supplementary Table 1. Overview of studies using tACS at frequencies outside of the theta or gamma range (not included in the meta-analysis) | | | | | | |
| --- | --- | --- | --- | --- | --- | --- |
| Reference | Montage | Freq. (Hz) | Intensity (mA) | Task | Time between encoding and retrieval | Hedges’ g |
| Luckey et al. 2022 | Bilateral C2 dermatome | 40 | 3 | word association task | 7 days | 0.17 |
| Shtoots et al., 2024 | Fz, AF4/AF3/FC2/FC1 | 16 | 1.5 | delayed recall, pictures | 2 hours | 0.06 |
|  | Fz, AF4/AF3/FC2/FC1 | 16 | 1.5 | delayed recall, pictures | 1 day | 0.08 |
|  | Fz, AF4/AF3/FC2/FC1 | 16 | 1.5 | delayed recall, pictures | 7 day | 0.08 |
| Shtoots et al., 2025 | Chin, P1/P2/P3/P4 | 16 | 1.5 | multiple choice exam, video | 20 min | 0.14 |
|  | Chin, P1/P2/P3/P4 | 16 | 1.5 | multiple choice exam, video | 1 day | 0.12 |
|  | Chin, P1/P2/P3/P4 | 16 | 1.5 | multiple choice exam, video | 7 day | 0.29 |
|  | P3, C3/TP7/Pz/O1 | 16 | 1.5 | multiple choice exam, video | 20 min | 0.16 |
|  | P3, C3/TP7/Pz/O1 | 16 | 1.5 | multiple choice exam, video | 1 day | 0.14 |
|  | P3, C3/TP7/Pz/O1 | 16 | 1.5 | multiple choice exam, video | 7 day | 0.17 |
|  | Fz, AF4/AF3/FC2/FC1 | 16 | 1.5 | multiple choice exam, video | 20 min | 0.10 |
|  | Fz, AF4/AF3/FC2/FC1 | 16 | 1.5 | multiple choice exam, video | 1 day | 0.09 |
|  | Fz, AF4/AF3/FC2/FC1 | 16 | 1.5 | multiple choice exam, video | 7 day | -0.12 |

| Supplementary Table 2. PEI values of theta tACS per brain region. The order in which the values are presented corresponds to the order of visual presentation in Figure 2. | | | | | | | | | |
| --- | --- | --- | --- | --- | --- | --- | --- | --- | --- |
| **Left Hemisphere** | | | | | **Right Hemisphere** | | | | |
| *Glasser*  *idx* | *name* | *Huang*  *idx* | *PEI* | *p* | *Glasser*  *idx* | *name* | *Huang*  *idx* | *PEI* | *p* |
| 1 | V1 | 1 | -0.073 | 0.672 | 181 | V1 | 1 | -0.045 | 0.794 |
| 6 | V4 | 2 | -0.133 | 0.439 | 185 | V3 | 2 | -0.060 | 0.730 |
| 5 | V3 | 2 | -0.122 | 0.479 | 184 | V2 | 2 | -0.059 | 0.733 |
| 4 | V2 | 2 | -0.097 | 0.572 | 186 | V4 | 2 | -0.048 | 0.781 |
| 17 | IPS1 | 3 | -0.174 | 0.311 | 332 | V6A | 3 | -0.122 | 0.477 |
| 152 | V6A | 3 | -0.167 | 0.330 | 197 | IPS1 | 3 | -0.120 | 0.486 |
| 16 | V7 | 3 | -0.147 | 0.393 | 196 | V7 | 3 | -0.107 | 0.534 |
| 19 | V3B | 3 | -0.144 | 0.403 | 193 | V3A | 3 | -0.102 | 0.554 |
| 13 | V3A | 3 | -0.123 | 0.473 | 199 | V3B | 3 | -0.077 | 0.654 |
| 3 | V6 | 3 | -0.100 | 0.560 | 183 | V6 | 3 | -0.071 | 0.680 |
| 163 | VVC | 4 | -0.334 | 0.046 | 340 | VMV2 | 4 | -0.137 | 0.424 |
| 154 | VMV3 | 4 | -0.316 | 0.060 | 333 | VMV1 | 4 | -0.130 | 0.451 |
| 18 | FFC | 4 | -0.295 | 0.080 | 334 | VMV3 | 4 | -0.119 | 0.489 |
| 160 | VMV2 | 4 | -0.287 | 0.090 | 343 | VVC | 4 | -0.111 | 0.520 |
| 153 | VMV1 | 4 | -0.276 | 0.103 | 198 | FFC | 4 | -0.101 | 0.560 |
| 7 | V8 | 4 | -0.265 | 0.118 | 187 | V8 | 4 | -0.077 | 0.655 |
| 22 | PIT | 4 | -0.125 | 0.468 | 202 | PIT | 4 | -0.018 | 0.915 |
| 138 | PH | 5 | -0.238 | 0.163 | 182 | MST | 5 | -0.082 | 0.632 |
| 2 | MST | 5 | -0.216 | 0.205 | 337 | FST | 5 | -0.076 | 0.659 |
| 157 | FST | 5 | -0.209 | 0.222 | 318 | PH | 5 | -0.074 | 0.670 |
| 23 | MT | 5 | -0.175 | 0.306 | 338 | V3CD | 5 | -0.067 | 0.697 |
| 159 | LO3 | 5 | -0.154 | 0.370 | 203 | MT | 5 | -0.059 | 0.734 |
| 156 | V4t | 5 | -0.144 | 0.403 | 339 | LO3 | 5 | -0.057 | 0.743 |
| 158 | V3CD | 5 | -0.143 | 0.406 | 200 | LO1 | 5 | -0.040 | 0.817 |
| 20 | LO1 | 5 | -0.125 | 0.466 | 336 | V4t | 5 | -0.037 | 0.832 |
| 21 | LO2 | 5 | -0.106 | 0.538 | 201 | LO2 | 5 | -0.017 | 0.922 |
| 53 | 3a | 6 | -0.437 | 0.008 | 233 | 3a | 6 | -0.250 | 0.142 |
| 9 | 3b | 6 | -0.387 | 0.019 | 188 | 4 | 6 | -0.230 | 0.177 |
| 8 | 4 | 6 | -0.385 | 0.020 | 189 | 3b | 6 | -0.227 | 0.184 |
| 51 | 1 | 6 | -0.351 | 0.035 | 231 | 1 | 6 | -0.200 | 0.242 |
| 52 | 2 | 6 | -0.326 | 0.052 | 232 | 2 | 6 | -0.198 | 0.247 |
| 55 | 6mp | 7 | -0.294 | 0.081 | 219 | 5L | 7 | -0.257 | 0.130 |
| 39 | 5L | 7 | -0.271 | 0.110 | 216 | 5m | 7 | -0.244 | 0.151 |
| 40 | 24dd | 7 | -0.264 | 0.119 | 220 | 24dd | 7 | -0.232 | 0.174 |
| 41 | 24dv | 7 | -0.255 | 0.133 | 235 | 6mp | 7 | -0.230 | 0.176 |
| 38 | 23c | 7 | -0.254 | 0.134 | 221 | 24dv | 7 | -0.230 | 0.177 |
| 37 | 5mv | 7 | -0.250 | 0.140 | 218 | 23c | 7 | -0.212 | 0.214 |
| 44 | 6ma | 7 | -0.240 | 0.158 | 217 | 5mv | 7 | -0.199 | 0.244 |
| 36 | 5m | 7 | -0.237 | 0.165 | 223 | SCEF | 7 | -0.184 | 0.281 |
| 43 | SCEF | 7 | -0.207 | 0.225 | 224 | 6ma | 7 | -0.170 | 0.322 |
| 78 | 6r | 8 | -0.545 | 0.001 | 258 | 6r | 8 | -0.243 | 0.154 |
| 56 | 6v | 8 | -0.509 | 0.001 | 276 | 6a | 8 | -0.237 | 0.164 |
| 11 | PEF | 8 | -0.473 | 0.004 | 190 | FEF | 8 | -0.226 | 0.185 |
| 10 | FEF | 8 | -0.447 | 0.006 | 191 | PEF | 8 | -0.220 | 0.198 |
| 12 | 55b | 8 | -0.441 | 0.007 | 236 | 6v | 8 | -0.215 | 0.209 |
| 96 | 6a | 8 | -0.381 | 0.022 | 234 | 6d | 8 | -0.202 | 0.236 |
| 54 | 6d | 8 | -0.339 | 0.043 | 192 | 55b | 8 | -0.202 | 0.237 |
| 99 | 43 | 9 | -0.521 | 0.001 | 279 | 43 | 9 | -0.207 | 0.225 |
| 113 | FOP1 | 9 | -0.498 | 0.002 | 293 | FOP1 | 9 | -0.199 | 0.245 |
| 100 | OP4 | 9 | -0.437 | 0.008 | 282 | OP2-3 | 9 | -0.178 | 0.298 |
| 102 | OP2-3 | 9 | -0.404 | 0.014 | 280 | OP4 | 9 | -0.172 | 0.317 |
| 101 | OP1 | 9 | -0.349 | 0.037 | 281 | OP1 | 9 | -0.140 | 0.416 |
| 103 | 52 | 10 | -0.440 | 0.007 | 283 | 52 | 10 | -0.214 | 0.210 |
| 173 | MBelt | 10 | -0.425 | 0.010 | 284 | RI | 10 | -0.170 | 0.320 |
| 124 | PBelt | 10 | -0.408 | 0.013 | 353 | MBelt | 10 | -0.169 | 0.323 |
| 174 | LBelt | 10 | -0.364 | 0.029 | 304 | PBelt | 10 | -0.164 | 0.340 |
| 104 | RI | 10 | -0.331 | 0.048 | 354 | LBelt | 10 | -0.132 | 0.444 |
| 24 | A1 | 10 | -0.299 | 0.076 | 285 | PFcm | 10 | -0.127 | 0.459 |
| 105 | PFcm | 10 | -0.279 | 0.099 | 204 | A1 | 10 | -0.106 | 0.536 |
| 128 | STSda | 11 | -0.521 | 0.001 | 287 | TA2 | 11 | -0.198 | 0.248 |
| 107 | TA2 | 11 | -0.511 | 0.001 | 303 | STGa | 11 | -0.193 | 0.259 |
| 125 | A5 | 11 | -0.500 | 0.002 | 308 | STSda | 11 | -0.183 | 0.285 |
| 176 | STSva | 11 | -0.491 | 0.002 | 355 | A4 | 11 | -0.180 | 0.292 |
| 123 | STGa | 11 | -0.489 | 0.002 | 305 | A5 | 11 | -0.176 | 0.306 |
| 175 | A4 | 11 | -0.433 | 0.008 | 356 | STSva | 11 | -0.162 | 0.345 |
| 129 | STSdp | 11 | -0.397 | 0.016 | 309 | STSdp | 11 | -0.152 | 0.377 |
| 130 | STSvp | 11 | -0.395 | 0.017 | 310 | STSvp | 11 | -0.137 | 0.424 |
| 108 | FOP4 | 12 | -0.485 | 0.003 | 290 | Pir | 12 | -0.254 | 0.134 |
| 114 | FOP3 | 12 | -0.464 | 0.004 | 292 | AAIC | 12 | -0.241 | 0.156 |
| 178 | PI | 12 | -0.463 | 0.004 | 349 | FOP5 | 12 | -0.238 | 0.162 |
| 109 | MI | 12 | -0.458 | 0.005 | 291 | AVI | 12 | -0.226 | 0.186 |
| 169 | FOP5 | 12 | -0.457 | 0.005 | 288 | FOP4 | 12 | -0.225 | 0.187 |
| 115 | FOP2 | 12 | -0.441 | 0.007 | 294 | FOP3 | 12 | -0.225 | 0.188 |
| 112 | AAIC | 12 | -0.436 | 0.008 | 347 | PoI1 | 12 | -0.208 | 0.223 |
| 111 | AVI | 12 | -0.432 | 0.008 | 289 | MI | 12 | -0.205 | 0.229 |
| 110 | Pir | 12 | -0.426 | 0.009 | 295 | FOP2 | 12 | -0.199 | 0.244 |
| 167 | PoI1 | 12 | -0.414 | 0.012 | 358 | PI | 12 | -0.192 | 0.263 |
| 106 | PoI2 | 12 | -0.397 | 0.016 | 286 | PoI2 | 12 | -0.161 | 0.347 |
| 168 | Ig | 12 | -0.376 | 0.024 | 348 | Ig | 12 | -0.158 | 0.357 |
| 135 | TF | 13 | -0.358 | 0.032 | 298 | EC | 13 | -0.188 | 0.272 |
| 120 | H | 13 | -0.340 | 0.042 | 300 | H | 13 | -0.163 | 0.343 |
| 127 | PHA3 | 13 | -0.329 | 0.050 | 299 | PreS | 13 | -0.144 | 0.401 |
| 118 | EC | 13 | -0.318 | 0.059 | 335 | PHA2 | 13 | -0.143 | 0.404 |
| 122 | PeEc | 13 | -0.300 | 0.075 | 315 | TF | 13 | -0.135 | 0.433 |
| 155 | PHA2 | 13 | -0.297 | 0.079 | 302 | PeEc | 13 | -0.135 | 0.434 |
| 126 | PHA1 | 13 | -0.264 | 0.120 | 307 | PHA3 | 13 | -0.133 | 0.439 |
| 119 | PreS | 13 | -0.248 | 0.145 | 306 | PHA1 | 13 | -0.120 | 0.484 |
| 132 | TE1a | 14 | -0.514 | 0.001 | 312 | TE1a | 14 | -0.177 | 0.302 |
| 177 | TE1m | 14 | -0.438 | 0.007 | 311 | TGd | 14 | -0.155 | 0.367 |
| 134 | TE2a | 14 | -0.394 | 0.017 | 357 | TE1m | 14 | -0.154 | 0.370 |
| 131 | TGd | 14 | -0.389 | 0.019 | 313 | TE1p | 14 | -0.148 | 0.390 |
| 136 | TE2p | 14 | -0.382 | 0.021 | 316 | TE2p | 14 | -0.141 | 0.414 |
| 133 | TE1p | 14 | -0.376 | 0.024 | 314 | TE2a | 14 | -0.140 | 0.415 |
| 172 | TGv | 14 | -0.348 | 0.038 | 352 | TGv | 14 | -0.133 | 0.439 |
| 137 | PHT | 14 | -0.286 | 0.090 | 317 | PHT | 14 | -0.117 | 0.497 |
| 139 | TPOJ1 | 15 | -0.304 | 0.071 | 205 | PSL | 15 | -0.154 | 0.368 |
| 28 | STV | 15 | -0.285 | 0.091 | 208 | STV | 15 | -0.131 | 0.448 |
| 25 | PSL | 15 | -0.267 | 0.116 | 319 | TPOJ1 | 15 | -0.118 | 0.494 |
| 140 | TPOJ2 | 15 | -0.251 | 0.139 | 320 | TPOJ2 | 15 | -0.108 | 0.531 |
| 141 | TPOJ3 | 15 | -0.194 | 0.258 | 321 | TPOJ3 | 15 | -0.085 | 0.620 |
| 47 | 7PC | 16 | -0.299 | 0.076 | 222 | 7AL | 16 | -0.231 | 0.176 |
| 42 | 7AL | 16 | -0.285 | 0.091 | 225 | 7Am | 16 | -0.228 | 0.180 |
| 117 | AIP | 16 | -0.277 | 0.102 | 229 | VIP | 16 | -0.207 | 0.225 |
| 49 | VIP | 16 | -0.257 | 0.130 | 227 | 7PC | 16 | -0.205 | 0.230 |
| 48 | LIPv | 16 | -0.250 | 0.141 | 226 | 7Pl | 16 | -0.194 | 0.257 |
| 45 | 7Am | 16 | -0.233 | 0.170 | 209 | 7Pm | 16 | -0.193 | 0.259 |
| 46 | 7Pl | 16 | -0.214 | 0.210 | 228 | LIPv | 16 | -0.174 | 0.309 |
| 95 | LIPd | 16 | -0.210 | 0.218 | 297 | AIP | 16 | -0.170 | 0.321 |
| 50 | MIP | 16 | -0.205 | 0.231 | 230 | MIP | 16 | -0.160 | 0.352 |
| 29 | 7Pm | 16 | -0.198 | 0.246 | 275 | LIPd | 16 | -0.159 | 0.353 |
| 147 | PFop | 17 | -0.368 | 0.027 | 327 | PFop | 17 | -0.192 | 0.262 |
| 116 | PFt | 17 | -0.331 | 0.048 | 296 | PFt | 17 | -0.182 | 0.288 |
| 144 | IP2 | 17 | -0.277 | 0.102 | 324 | IP2 | 17 | -0.170 | 0.322 |
| 148 | PF | 17 | -0.270 | 0.111 | 328 | PF | 17 | -0.160 | 0.351 |
| 149 | PFm | 17 | -0.244 | 0.151 | 329 | PFm | 17 | -0.158 | 0.356 |
| 150 | PGi | 17 | -0.223 | 0.192 | 325 | IP1 | 17 | -0.146 | 0.395 |
| 145 | IP1 | 17 | -0.207 | 0.226 | 331 | PGs | 17 | -0.119 | 0.490 |
| 146 | IP0 | 17 | -0.190 | 0.267 | 326 | IP0 | 17 | -0.108 | 0.532 |
| 151 | PGs | 17 | -0.167 | 0.330 | 330 | PGi | 17 | -0.105 | 0.542 |
| 143 | PGp | 17 | -0.155 | 0.367 | 323 | PGp | 17 | -0.078 | 0.649 |
| 162 | 31a | 18 | -0.246 | 0.147 | 342 | 31a | 18 | -0.229 | 0.180 |
| 161 | 31pd | 18 | -0.230 | 0.178 | 212 | 23d | 18 | -0.206 | 0.227 |
| 35 | 31pv | 18 | -0.218 | 0.201 | 207 | PCV | 18 | -0.201 | 0.238 |
| 27 | PCV | 18 | -0.217 | 0.204 | 215 | 31pv | 18 | -0.198 | 0.247 |
| 32 | 23d | 18 | -0.214 | 0.209 | 341 | 31pd | 18 | -0.193 | 0.260 |
| 121 | ProS | 18 | -0.208 | 0.224 | 194 | RSC | 18 | -0.178 | 0.300 |
| 14 | RSC | 18 | -0.201 | 0.238 | 214 | d23ab | 18 | -0.167 | 0.330 |
| 34 | d23ab | 18 | -0.170 | 0.322 | 210 | 7m | 18 | -0.140 | 0.414 |
| 30 | 7m | 18 | -0.149 | 0.385 | 301 | ProS | 18 | -0.133 | 0.439 |
| 15 | POS2 | 18 | -0.136 | 0.428 | 213 | v23ab | 18 | -0.124 | 0.472 |
| 142 | DVT | 18 | -0.126 | 0.464 | 195 | POS2 | 18 | -0.121 | 0.481 |
| 33 | v23ab | 18 | -0.123 | 0.475 | 322 | DVT | 18 | -0.101 | 0.560 |
| 31 | POS1 | 18 | -0.106 | 0.538 | 211 | POS1 | 18 | -0.081 | 0.638 |
| 166 | pOFC | 19 | -0.340 | 0.042 | 344 | 25 | 19 | -0.254 | 0.135 |
| 164 | 25 | 19 | -0.259 | 0.127 | 346 | pOFC | 19 | -0.248 | 0.144 |
| 64 | p32 | 19 | -0.246 | 0.148 | 268 | 10v | 19 | -0.220 | 0.197 |
| 88 | 10v | 19 | -0.242 | 0.155 | 345 | s32 | 19 | -0.216 | 0.205 |
| 165 | s32 | 19 | -0.240 | 0.158 | 241 | a24 | 19 | -0.200 | 0.242 |
| 61 | a24 | 19 | -0.216 | 0.206 | 237 | p24pr | 19 | -0.198 | 0.247 |
| 65 | 10r | 19 | -0.208 | 0.223 | 238 | 33pr | 19 | -0.186 | 0.278 |
| 57 | p24pr | 19 | -0.206 | 0.227 | 244 | p32 | 19 | -0.182 | 0.287 |
| 60 | p32pr | 19 | -0.204 | 0.233 | 245 | 10r | 19 | -0.172 | 0.315 |
| 58 | 33pr | 19 | -0.195 | 0.255 | 240 | p32pr | 19 | -0.163 | 0.343 |
| 179 | a32pr | 19 | -0.167 | 0.330 | 239 | a24pr | 19 | -0.152 | 0.377 |
| 59 | a24pr | 19 | -0.167 | 0.331 | 359 | a32pr | 19 | -0.116 | 0.502 |
| 180 | p24 | 19 | -0.138 | 0.423 | 360 | p24 | 19 | -0.104 | 0.545 |
| 62 | d32 | 19 | -0.115 | 0.503 | 242 | d32 | 19 | -0.058 | 0.735 |
| 63 | 8BM | 19 | -0.032 | 0.855 | 243 | 8BM | 19 | -0.017 | 0.920 |
| 69 | 9m | 19 | -0.003 | 0.987 | 249 | 9m | 19 | 0.038 | 0.824 |
| 94 | 47s | 20 | -0.377 | 0.023 | 270 | 10pp | 20 | -0.261 | 0.124 |
| 66 | 47m | 20 | -0.375 | 0.024 | 273 | OFC | 20 | -0.250 | 0.140 |
| 91 | 11l | 20 | -0.372 | 0.025 | 271 | 11l | 20 | -0.224 | 0.188 |
| 89 | a10p | 20 | -0.346 | 0.038 | 269 | a10p | 20 | -0.206 | 0.228 |
| 92 | 13l | 20 | -0.341 | 0.042 | 272 | 13l | 20 | -0.205 | 0.229 |
| 90 | 10pp | 20 | -0.335 | 0.046 | 246 | 47m | 20 | -0.196 | 0.251 |
| 93 | OFC | 20 | -0.309 | 0.067 | 274 | 47s | 20 | -0.188 | 0.273 |
| 170 | p10p | 20 | -0.243 | 0.153 | 252 | 10d | 20 | -0.081 | 0.640 |
| 72 | 10d | 20 | -0.151 | 0.380 | 350 | p10p | 20 | -0.077 | 0.653 |
| 74 | 44 | 21 | -0.526 | 0.001 | 260 | IFJp | 21 | -0.264 | 0.119 |
| 79 | IFJa | 21 | -0.503 | 0.002 | 254 | 44 | 21 | -0.227 | 0.182 |
| 80 | IFJp | 21 | -0.496 | 0.002 | 255 | 45 | 21 | -0.221 | 0.195 |
| 75 | 45 | 21 | -0.487 | 0.003 | 259 | IFJa | 21 | -0.217 | 0.204 |
| 81 | IFSp | 21 | -0.475 | 0.003 | 256 | 47l | 21 | -0.199 | 0.245 |
| 82 | IFSa | 21 | -0.440 | 0.007 | 257 | a47r | 21 | -0.189 | 0.269 |
| 76 | 47l | 21 | -0.425 | 0.010 | 351 | p47r | 21 | -0.179 | 0.296 |
| 77 | a47r | 21 | -0.422 | 0.010 | 262 | IFSa | 21 | -0.174 | 0.310 |
| 171 | p47r | 21 | -0.415 | 0.012 | 261 | IFSp | 21 | -0.174 | 0.311 |
| 73 | 8C | 22 | -0.462 | 0.004 | 253 | 8C | 22 | -0.201 | 0.239 |
| 83 | p9-46v | 22 | -0.422 | 0.010 | 277 | i6-8 | 22 | -0.171 | 0.319 |
| 67 | 8Av | 22 | -0.376 | 0.024 | 247 | 8Av | 22 | -0.170 | 0.321 |
| 97 | i6-8 | 22 | -0.343 | 0.041 | 263 | p9-46v | 22 | -0.136 | 0.429 |
| 85 | a9-46v | 22 | -0.334 | 0.046 | 265 | a9-46v | 22 | -0.101 | 0.557 |
| 84 | 46 | 22 | -0.309 | 0.067 | 264 | 46 | 22 | -0.074 | 0.669 |
| 86 | 9-46d | 22 | -0.235 | 0.168 | 206 | SFL | 22 | -0.072 | 0.677 |
| 68 | 8Ad | 22 | -0.217 | 0.204 | 278 | s6-8 | 22 | -0.055 | 0.750 |
| 98 | s6-8 | 22 | -0.150 | 0.381 | 248 | 8Ad | 22 | -0.045 | 0.795 |
| 26 | SFL | 22 | -0.110 | 0.524 | 266 | 9-46d | 22 | -0.002 | 0.991 |
| 87 | 9a | 22 | -0.065 | 0.706 | 267 | 9a | 22 | 0.102 | 0.554 |
| 71 | 9p | 22 | -0.017 | 0.924 | 251 | 9p | 22 | 0.108 | 0.531 |
| 70 | 8BL | 22 | 0.052 | 0.761 | 250 | 8BL | 22 | 0.144 | 0.403 |
| Supplementary Table 3. PEI values for gamma tACS per brain region. The order in which the values are presented corresponds to the order of visual presentation in Figure 3. | | | | | | | | | |
| **Left Hemisphere** | | | | | **Right Hemisphere** | | | | |
| *Glasser*  *idx* | *name* | *Huang*  *idx* | *PEI* | *p* | *Glasser*  *idx* | *name* | *Huang*  *idx* | *PEI* | *p* |
| 1 | V1 | 1 | 0.514 | 0.014 | 181 | V1 | 1 | 0.473 | 0.026 |
| 6 | V4 | 2 | 0.497 | 0.018 | 186 | V4 | 2 | 0.434 | 0.043 |
| 5 | V3 | 2 | 0.500 | 0.017 | 185 | V3 | 2 | 0.460 | 0.031 |
| 4 | V2 | 2 | 0.502 | 0.017 | 184 | V2 | 2 | 0.476 | 0.025 |
| 17 | IPS1 | 3 | 0.456 | 0.033 | 197 | IPS1 | 3 | 0.345 | 0.115 |
| 152 | V6A | 3 | 0.461 | 0.030 | 332 | V6A | 3 | 0.391 | 0.071 |
| 19 | V3B | 3 | 0.480 | 0.023 | 196 | V7 | 3 | 0.395 | 0.068 |
| 16 | V7 | 3 | 0.482 | 0.023 | 199 | V3B | 3 | 0.398 | 0.066 |
| 3 | V6 | 3 | 0.497 | 0.018 | 183 | V6 | 3 | 0.429 | 0.045 |
| 13 | V3A | 3 | 0.504 | 0.016 | 193 | V3A | 3 | 0.432 | 0.044 |
| 163 | VVC | 4 | 0.187 | 0.404 | 198 | FFC | 4 | 0.199 | 0.375 |
| 153 | VMV1 | 4 | 0.190 | 0.396 | 340 | VMV2 | 4 | 0.252 | 0.258 |
| 154 | VMV3 | 4 | 0.278 | 0.210 | 343 | VVC | 4 | 0.264 | 0.235 |
| 160 | VMV2 | 4 | 0.285 | 0.198 | 334 | VMV3 | 4 | 0.299 | 0.176 |
| 18 | FFC | 4 | 0.306 | 0.166 | 333 | VMV1 | 4 | 0.320 | 0.146 |
| 7 | V8 | 4 | 0.371 | 0.089 | 187 | V8 | 4 | 0.361 | 0.098 |
| 22 | PIT | 4 | 0.493 | 0.019 | 202 | PIT | 4 | 0.393 | 0.070 |
| 157 | FST | 5 | 0.355 | 0.104 | 318 | PH | 5 | 0.213 | 0.340 |
| 138 | PH | 5 | 0.359 | 0.100 | 337 | FST | 5 | 0.219 | 0.327 |
| 2 | MST | 5 | 0.373 | 0.086 | 182 | MST | 5 | 0.244 | 0.273 |
| 23 | MT | 5 | 0.384 | 0.077 | 203 | MT | 5 | 0.297 | 0.179 |
| 159 | LO3 | 5 | 0.384 | 0.077 | 339 | LO3 | 5 | 0.325 | 0.139 |
| 156 | V4t | 5 | 0.453 | 0.034 | 336 | V4t | 5 | 0.344 | 0.116 |
| 20 | LO1 | 5 | 0.460 | 0.031 | 200 | LO1 | 5 | 0.394 | 0.069 |
| 158 | V3CD | 5 | 0.462 | 0.030 | 201 | LO2 | 5 | 0.398 | 0.066 |
| 21 | LO2 | 5 | 0.493 | 0.019 | 338 | V3CD | 5 | 0.422 | 0.050 |
| 53 | 3a | 6 | 0.067 | 0.766 | 233 | 3a | 6 | 0.012 | 0.957 |
| 9 | 3b | 6 | 0.078 | 0.731 | 188 | 4 | 6 | 0.028 | 0.903 |
| 8 | 4 | 6 | 0.087 | 0.701 | 231 | 1 | 6 | 0.033 | 0.885 |
| 51 | 1 | 6 | 0.100 | 0.659 | 189 | 3b | 6 | 0.041 | 0.857 |
| 52 | 2 | 6 | 0.139 | 0.536 | 232 | 2 | 6 | 0.089 | 0.693 |
| 41 | 24dv | 7 | 0.030 | 0.894 | 224 | 6ma | 7 | 0.010 | 0.965 |
| 43 | SCEF | 7 | 0.055 | 0.806 | 223 | SCEF | 7 | 0.011 | 0.959 |
| 44 | 6ma | 7 | 0.060 | 0.792 | 221 | 24dv | 7 | 0.038 | 0.868 |
| 40 | 24dd | 7 | 0.107 | 0.634 | 235 | 6mp | 7 | 0.057 | 0.802 |
| 55 | 6mp | 7 | 0.140 | 0.535 | 220 | 24dd | 7 | 0.113 | 0.615 |
| 38 | 23c | 7 | 0.180 | 0.423 | 219 | 5L | 7 | 0.163 | 0.468 |
| 37 | 5mv | 7 | 0.262 | 0.238 | 216 | 5m | 7 | 0.165 | 0.464 |
| 36 | 5m | 7 | 0.282 | 0.203 | 217 | 5mv | 7 | 0.207 | 0.354 |
| 39 | 5L | 7 | 0.324 | 0.141 | 218 | 23c | 7 | 0.215 | 0.335 |
| 78 | 6r | 8 | -0.063 | 0.782 | 258 | 6r | 8 | -0.145 | 0.519 |
| 56 | 6v | 8 | -0.021 | 0.925 | 236 | 6v | 8 | -0.132 | 0.557 |
| 12 | 55b | 8 | 0.007 | 0.975 | 191 | PEF | 8 | -0.086 | 0.702 |
| 10 | FEF | 8 | 0.010 | 0.964 | 192 | 55b | 8 | -0.049 | 0.830 |
| 11 | PEF | 8 | 0.018 | 0.935 | 276 | 6a | 8 | -0.035 | 0.878 |
| 96 | 6a | 8 | 0.044 | 0.847 | 190 | FEF | 8 | -0.027 | 0.904 |
| 54 | 6d | 8 | 0.116 | 0.607 | 234 | 6d | 8 | 0.030 | 0.895 |
| 99 | 43 | 9 | -0.033 | 0.883 | 279 | 43 | 9 | -0.156 | 0.487 |
| 113 | FOP1 | 9 | -0.027 | 0.905 | 293 | FOP1 | 9 | -0.145 | 0.520 |
| 100 | OP4 | 9 | -0.002 | 0.993 | 280 | OP4 | 9 | -0.099 | 0.660 |
| 102 | OP2-3 | 9 | 0.037 | 0.869 | 281 | OP1 | 9 | -0.003 | 0.989 |
| 101 | OP1 | 9 | 0.102 | 0.652 | 282 | OP2-3 | 9 | -0.002 | 0.993 |
| 173 | MBelt | 10 | -0.005 | 0.981 | 283 | 52 | 10 | -0.088 | 0.696 |
| 124 | PBelt | 10 | 0.004 | 0.987 | 304 | PBelt | 10 | -0.080 | 0.724 |
| 103 | 52 | 10 | 0.013 | 0.955 | 353 | MBelt | 10 | -0.069 | 0.760 |
| 174 | LBelt | 10 | 0.084 | 0.709 | 354 | LBelt | 10 | -0.031 | 0.892 |
| 104 | RI | 10 | 0.110 | 0.626 | 204 | A1 | 10 | -0.027 | 0.905 |
| 24 | A1 | 10 | 0.117 | 0.602 | 284 | RI | 10 | -0.003 | 0.988 |
| 105 | PFcm | 10 | 0.124 | 0.581 | 285 | PFcm | 10 | 0.016 | 0.942 |
| 123 | STGa | 11 | -0.136 | 0.545 | 287 | TA2 | 11 | -0.095 | 0.673 |
| 128 | STSda | 11 | -0.125 | 0.578 | 355 | A4 | 11 | -0.081 | 0.720 |
| 107 | TA2 | 11 | -0.114 | 0.614 | 305 | A5 | 11 | -0.078 | 0.728 |
| 125 | A5 | 11 | -0.106 | 0.640 | 308 | STSda | 11 | -0.067 | 0.767 |
| 176 | STSva | 11 | -0.104 | 0.646 | 356 | STSva | 11 | -0.066 | 0.771 |
| 175 | A4 | 11 | -0.070 | 0.758 | 309 | STSdp | 11 | -0.033 | 0.884 |
| 130 | STSvp | 11 | 0.007 | 0.976 | 303 | STGa | 11 | -0.032 | 0.887 |
| 129 | STSdp | 11 | 0.013 | 0.955 | 310 | STSvp | 11 | -0.032 | 0.887 |
| 169 | FOP5 | 12 | -0.129 | 0.567 | 349 | FOP5 | 12 | -0.142 | 0.529 |
| 111 | AVI | 12 | -0.106 | 0.637 | 347 | PoI1 | 12 | -0.132 | 0.556 |
| 108 | FOP4 | 12 | -0.091 | 0.688 | 290 | Pir | 12 | -0.123 | 0.585 |
| 178 | PI | 12 | -0.088 | 0.696 | 291 | AVI | 12 | -0.110 | 0.626 |
| 109 | MI | 12 | -0.076 | 0.737 | 292 | AAIC | 12 | -0.103 | 0.649 |
| 114 | FOP3 | 12 | -0.063 | 0.781 | 288 | FOP4 | 12 | -0.102 | 0.653 |
| 112 | AAIC | 12 | -0.049 | 0.829 | 286 | PoI2 | 12 | -0.095 | 0.673 |
| 110 | Pir | 12 | -0.041 | 0.857 | 289 | MI | 12 | -0.095 | 0.674 |
| 106 | PoI2 | 12 | 0.009 | 0.968 | 294 | FOP3 | 12 | -0.079 | 0.725 |
| 115 | FOP2 | 12 | 0.018 | 0.935 | 295 | FOP2 | 12 | -0.058 | 0.797 |
| 167 | PoI1 | 12 | 0.024 | 0.916 | 358 | PI | 12 | -0.045 | 0.843 |
| 168 | Ig | 12 | 0.052 | 0.819 | 348 | Ig | 12 | -0.035 | 0.876 |
| 122 | PeEc | 13 | -0.061 | 0.786 | 315 | TF | 13 | 0.003 | 0.990 |
| 118 | EC | 13 | -0.059 | 0.795 | 300 | H | 13 | 0.078 | 0.730 |
| 135 | TF | 13 | -0.030 | 0.895 | 298 | EC | 13 | 0.089 | 0.694 |
| 120 | H | 13 | -0.015 | 0.947 | 302 | PeEc | 13 | 0.101 | 0.656 |
| 126 | PHA1 | 13 | 0.080 | 0.725 | 335 | PHA2 | 13 | 0.142 | 0.529 |
| 119 | PreS | 13 | 0.088 | 0.697 | 307 | PHA3 | 13 | 0.149 | 0.507 |
| 127 | PHA3 | 13 | 0.100 | 0.658 | 299 | PreS | 13 | 0.149 | 0.507 |
| 155 | PHA2 | 13 | 0.104 | 0.644 | 306 | PHA1 | 13 | 0.240 | 0.281 |
| 132 | TE1a | 14 | -0.143 | 0.526 | 357 | TE1m | 14 | -0.085 | 0.707 |
| 131 | TGd | 14 | -0.102 | 0.652 | 312 | TE1a | 14 | -0.084 | 0.709 |
| 177 | TE1m | 14 | -0.093 | 0.680 | 314 | TE2a | 14 | -0.078 | 0.730 |
| 172 | TGv | 14 | -0.068 | 0.765 | 313 | TE1p | 14 | -0.046 | 0.839 |
| 134 | TE2a | 14 | -0.063 | 0.781 | 316 | TE2p | 14 | 0.004 | 0.987 |
| 133 | TE1p | 14 | 0.046 | 0.837 | 352 | TGv | 14 | 0.043 | 0.850 |
| 136 | TE2p | 14 | 0.094 | 0.676 | 311 | TGd | 14 | 0.047 | 0.837 |
| 137 | PHT | 14 | 0.156 | 0.487 | 317 | PHT | 14 | 0.049 | 0.828 |
| 28 | STV | 15 | 0.051 | 0.821 | 205 | PSL | 15 | 0.008 | 0.972 |
| 25 | PSL | 15 | 0.082 | 0.716 | 208 | STV | 15 | 0.042 | 0.854 |
| 139 | TPOJ1 | 15 | 0.098 | 0.666 | 319 | TPOJ1 | 15 | 0.054 | 0.811 |
| 140 | TPOJ2 | 15 | 0.218 | 0.328 | 320 | TPOJ2 | 15 | 0.117 | 0.603 |
| 141 | TPOJ3 | 15 | 0.332 | 0.130 | 321 | TPOJ3 | 15 | 0.253 | 0.254 |
| 47 | 7PC | 16 | 0.193 | 0.390 | 297 | AIP | 16 | 0.113 | 0.616 |
| 117 | AIP | 16 | 0.209 | 0.350 | 227 | 7PC | 16 | 0.183 | 0.414 |
| 48 | LIPv | 16 | 0.281 | 0.204 | 275 | LIPd | 16 | 0.188 | 0.400 |
| 42 | 7AL | 16 | 0.304 | 0.169 | 222 | 7AL | 16 | 0.195 | 0.384 |
| 49 | VIP | 16 | 0.328 | 0.135 | 228 | LIPv | 16 | 0.219 | 0.326 |
| 95 | LIPd | 16 | 0.342 | 0.119 | 225 | 7Am | 16 | 0.258 | 0.246 |
| 45 | 7Am | 16 | 0.375 | 0.085 | 229 | VIP | 16 | 0.260 | 0.242 |
| 50 | MIP | 16 | 0.405 | 0.061 | 230 | MIP | 16 | 0.287 | 0.194 |
| 29 | 7Pm | 16 | 0.414 | 0.055 | 226 | 7Pl | 16 | 0.313 | 0.155 |
| 46 | 7Pl | 16 | 0.416 | 0.053 | 209 | 7Pm | 16 | 0.355 | 0.104 |
| 147 | PFop | 17 | -0.005 | 0.981 | 327 | PFop | 17 | -0.061 | 0.787 |
| 116 | PFt | 17 | 0.022 | 0.924 | 296 | PFt | 17 | 0.015 | 0.946 |
| 148 | PF | 17 | 0.047 | 0.837 | 328 | PF | 17 | 0.024 | 0.916 |
| 149 | PFm | 17 | 0.162 | 0.470 | 329 | PFm | 17 | 0.135 | 0.550 |
| 144 | IP2 | 17 | 0.202 | 0.367 | 324 | IP2 | 17 | 0.149 | 0.508 |
| 150 | PGi | 17 | 0.251 | 0.259 | 330 | PGi | 17 | 0.184 | 0.411 |
| 151 | PGs | 17 | 0.315 | 0.152 | 325 | IP1 | 17 | 0.263 | 0.236 |
| 145 | IP1 | 17 | 0.366 | 0.093 | 331 | PGs | 17 | 0.270 | 0.224 |
| 143 | PGp | 17 | 0.399 | 0.065 | 323 | PGp | 17 | 0.348 | 0.112 |
| 146 | IP0 | 17 | 0.419 | 0.051 | 326 | IP0 | 17 | 0.362 | 0.097 |
| 32 | 23d | 18 | 0.148 | 0.511 | 212 | 23d | 18 | 0.150 | 0.504 |
| 162 | 31a | 18 | 0.202 | 0.366 | 301 | ProS | 18 | 0.209 | 0.350 |
| 14 | RSC | 18 | 0.235 | 0.291 | 342 | 31a | 18 | 0.219 | 0.326 |
| 34 | d23ab | 18 | 0.268 | 0.227 | 214 | d23ab | 18 | 0.246 | 0.269 |
| 121 | ProS | 18 | 0.269 | 0.226 | 194 | RSC | 18 | 0.252 | 0.257 |
| 35 | 31pv | 18 | 0.297 | 0.178 | 215 | 31pv | 18 | 0.278 | 0.209 |
| 27 | PCV | 18 | 0.320 | 0.146 | 207 | PCV | 18 | 0.284 | 0.199 |
| 161 | 31pd | 18 | 0.345 | 0.115 | 341 | 31pd | 18 | 0.309 | 0.162 |
| 33 | v23ab | 18 | 0.396 | 0.067 | 211 | POS1 | 18 | 0.340 | 0.121 |
| 31 | POS1 | 18 | 0.408 | 0.059 | 322 | DVT | 18 | 0.341 | 0.120 |
| 30 | 7m | 18 | 0.410 | 0.058 | 213 | v23ab | 18 | 0.343 | 0.118 |
| 142 | DVT | 18 | 0.453 | 0.034 | 210 | 7m | 18 | 0.401 | 0.063 |
| 15 | POS2 | 18 | 0.475 | 0.025 | 195 | POS2 | 18 | 0.408 | 0.059 |
| 164 | 25 | 19 | -0.185 | 0.409 | 249 | 9m | 19 | -0.152 | 0.498 |
| 88 | 10v | 19 | -0.178 | 0.429 | 242 | d32 | 19 | -0.105 | 0.641 |
| 64 | p32 | 19 | -0.173 | 0.441 | 268 | 10v | 19 | -0.103 | 0.647 |
| 65 | 10r | 19 | -0.165 | 0.463 | 244 | p32 | 19 | -0.101 | 0.654 |
| 166 | pOFC | 19 | -0.162 | 0.472 | 245 | 10r | 19 | -0.087 | 0.700 |
| 165 | s32 | 19 | -0.161 | 0.474 | 241 | a24 | 19 | -0.085 | 0.707 |
| 61 | a24 | 19 | -0.139 | 0.536 | 360 | p24 | 19 | -0.082 | 0.718 |
| 69 | 9m | 19 | -0.136 | 0.546 | 344 | 25 | 19 | -0.073 | 0.748 |
| 62 | d32 | 19 | -0.112 | 0.620 | 359 | a32pr | 19 | -0.069 | 0.760 |
| 179 | a32pr | 19 | -0.092 | 0.683 | 345 | s32 | 19 | -0.066 | 0.772 |
| 180 | p24 | 19 | -0.078 | 0.728 | 243 | 8BM | 19 | -0.056 | 0.805 |
| 63 | 8BM | 19 | -0.039 | 0.863 | 346 | pOFC | 19 | -0.053 | 0.816 |
| 60 | p32pr | 19 | -0.028 | 0.900 | 239 | a24pr | 19 | -0.024 | 0.917 |
| 59 | a24pr | 19 | -0.020 | 0.929 | 240 | p32pr | 19 | -0.010 | 0.963 |
| 58 | 33pr | 19 | -0.010 | 0.964 | 238 | 33pr | 19 | 0.010 | 0.964 |
| 57 | p24pr | 19 | 0.060 | 0.790 | 237 | p24pr | 19 | 0.058 | 0.798 |
| 93 | OFC | 20 | -0.206 | 0.356 | 252 | 10d | 20 | -0.199 | 0.375 |
| 170 | p10p | 20 | -0.206 | 0.357 | 246 | 47m | 20 | -0.157 | 0.486 |
| 89 | a10p | 20 | -0.193 | 0.389 | 350 | p10p | 20 | -0.145 | 0.518 |
| 72 | 10d | 20 | -0.187 | 0.404 | 270 | 10pp | 20 | -0.134 | 0.552 |
| 90 | 10pp | 20 | -0.177 | 0.430 | 269 | a10p | 20 | -0.132 | 0.557 |
| 91 | 11l | 20 | -0.167 | 0.457 | 271 | 11l | 20 | -0.106 | 0.640 |
| 92 | 13l | 20 | -0.139 | 0.536 | 274 | 47s | 20 | -0.091 | 0.687 |
| 66 | 47m | 20 | -0.107 | 0.637 | 272 | 13l | 20 | -0.084 | 0.711 |
| 94 | 47s | 20 | -0.071 | 0.754 | 273 | OFC | 20 | -0.065 | 0.774 |
| 171 | p47r | 21 | -0.171 | 0.446 | 255 | 45 | 21 | -0.220 | 0.324 |
| 77 | a47r | 21 | -0.165 | 0.464 | 256 | 47l | 21 | -0.209 | 0.351 |
| 82 | IFSa | 21 | -0.124 | 0.583 | 262 | IFSa | 21 | -0.174 | 0.439 |
| 76 | 47l | 21 | -0.088 | 0.697 | 254 | 44 | 21 | -0.171 | 0.446 |
| 75 | 45 | 21 | -0.087 | 0.702 | 261 | IFSp | 21 | -0.162 | 0.470 |
| 81 | IFSp | 21 | -0.062 | 0.784 | 351 | p47r | 21 | -0.154 | 0.492 |
| 74 | 44 | 21 | -0.058 | 0.796 | 257 | a47r | 21 | -0.154 | 0.492 |
| 79 | IFJa | 21 | -0.058 | 0.798 | 259 | IFJa | 21 | -0.141 | 0.532 |
| 80 | IFJp | 21 | -0.033 | 0.884 | 260 | IFJp | 21 | -0.121 | 0.591 |
| 85 | a9-46v | 22 | -0.232 | 0.297 | 263 | p9-46v | 22 | -0.150 | 0.505 |
| 86 | 9-46d | 22 | -0.198 | 0.376 | 265 | a9-46v | 22 | -0.141 | 0.532 |
| 87 | 9a | 22 | -0.196 | 0.383 | 267 | 9a | 22 | -0.130 | 0.563 |
| 84 | 46 | 22 | -0.179 | 0.424 | 266 | 9-46d | 22 | -0.128 | 0.569 |
| 71 | 9p | 22 | -0.134 | 0.551 | 264 | 46 | 22 | -0.128 | 0.570 |
| 83 | p9-46v | 22 | -0.112 | 0.620 | 253 | 8C | 22 | -0.122 | 0.587 |
| 73 | 8C | 22 | -0.062 | 0.784 | 251 | 9p | 22 | -0.109 | 0.631 |
| 70 | 8BL | 22 | -0.047 | 0.836 | 248 | 8Ad | 22 | -0.100 | 0.658 |
| 68 | 8Ad | 22 | -0.046 | 0.839 | 247 | 8Av | 22 | -0.086 | 0.702 |
| 67 | 8Av | 22 | -0.045 | 0.842 | 250 | 8BL | 22 | -0.084 | 0.709 |
| 98 | s6-8 | 22 | -0.014 | 0.951 | 278 | s6-8 | 22 | -0.054 | 0.812 |
| 97 | i6-8 | 22 | 0.007 | 0.976 | 277 | i6-8 | 22 | -0.052 | 0.818 |
| 26 | SFL | 22 | 0.032 | 0.889 | 206 | SFL | 22 | -0.031 | 0.889 |
